# Diversity, host-specificity, and environmental drivers of the coral reef eukaryome

**DOI:** 10.64898/2026.06.02.729273

**Authors:** Laura del Rio-Hortega, Nicolas Henry, Clément Leboine, Christian R. Voolstra, Benjamin C.C. Hume, Kim-Isabelle Mayer, Pedro C. Junger, Clémentine Moulin, Emilie Boissin, Guillaume Bourdin, Guillaume Iwankow, Julie Poulain, Sarah Romac, Sylvain Agostini, Chris Bowler, Colomban de Vargas, Eric Douville, J. Michel Flores, Didier Forcioli, Paola Furla, Pierre E. Galand, Eric Gilson, Fabien Lombard, David A. Paz-Garcia, Stéphane Pesant, Stéphanie Reynaud, Shinichi Sunagawa, Olivier P. Thomas, Maggie M. Reddy, Romain Troublé, Rebecca Vega Thurber, Patrick Wincker, Didier Zoccola, Denis Allemand, Serge Planes, Betina M. Porcel, Quentin Carradec, Maren Ziegler

## Abstract

Coral reefs are among the most diverse ecosystems on Earth, yet the diversity and structure of their associated microeukaryotic communities remain poorly resolved. We characterized reef-associated eukaryomes across 113 reefs spanning the Pacific Ocean using more than 6,300 samples from corals, seawater, and sediments during the *Tara* Pacific expedition. We identified ∼121,000 eukaryotic ASVs, revealing one of Earth’s largest undocumented reservoirs of eukaryotic diversity; over 80% of the recovered diversity was previously undetected in global ocean surveys and fewer than 2% of sequences matched reference databases. Reef habitats supported highly distinct communities, with sediments and seawater harboring 20–40-fold higher richness than corals. Unexpectedly, coral-associated communities across 29 host lineages were consistently dominated by small metazoans, particularly demosponges and maxillopods, identifying these taxa as pervasive and previously unrecognized components of coral eukaryomes alongside Apicomplexa. Across the Pacific, eukaryome composition was strongly structured by environmental gradients, with thermal stress emerging as the primary driver of community turnover. Together, these results identify coral reefs as a globally important reservoir of hidden eukaryotic diversity and reveal the reef eukaryome as a sensitive indicator of ecosystem reorganization under climate change.

## INTRODUCTION

Coral reefs represent one of the most diverse ecosystems on Earth supporting the livelihoods of over 500 million people (*1–3*). They are now crossing critical ecological tipping points, with devastating and cascading consequences for all associated life (*4–6*). To anticipate and mitigate these impacts, it is essential to first understand the full extent of reef biodiversity, including its hidden microbial dimensions that support coral health and ecosystem functioning (*7–9*).

Both prokaryotic and eukaryotic microorganisms are integral to reef ecosystems, occurring as free-living forms and engaging in diverse symbiotic relationships within and across all kingdoms of life (*10*, *11*). The community of host-associated microeukaryotes, encompassing protists, fungi, and other microscopic eukaryotic organisms, has been termed the "eukaryome", a term proposed to recognise that microbial communities associated with a host are not composed exclusively of prokaryotes (*12*, *13*). Yet, despite their ecological importance (*14–17*), research on the coral reef microbiome has mainly focused on bacteria (*7*, *9*, *18–21*), leaving coral reef eukaryome poorly characterised.

Existing studies on the coral eukaryome have been mainly focused on the photoendosymbiotic dinoflagellate Symbiodiniaceae, which are crucial for coral survival (*22–24*), as well as disease-related protists and parasites (*25*, *26*). The application of metabarcoding techniques has revealed other prevalent and diverse coral associated eukaryotes, including hydrozoans, the chlorophyte *Ostreobium* and apicomplexans (*27–31*). Few studies to date have described the coral eukaryome, but these remain limited in geographic coverage or replication (*14*, *16*, *32*).

The effect of the biotic and abiotic environment on the eukaryome is also particularly important, yet largely unexplored. Environmental conditions and host traits are known to shape prokaryotic communities (*33–36*). However, it remains unclear whether coral eukaryomes primarily reflect the local environment or maintain a host-specific signature. Addressing these questions requires a large-scale, standardised survey that includes different coral hosts, habitats, and environmental gradients, an effort made possible through the *Tara* Pacific Expedition (*37*).

The *Tara* Pacific Expedition sought to place coral reefs among the first ecosystems with a comprehensive description of their large-scale genomic diversity (*37–39*). In this study, we aimed to i) characterise the eukaryome of coral holobionts and their surrounding environment (seawater and sediments) throughout the Pacific reefs, as well as the open ocean between them, ii) assess the habitat partitioning of these eukaryotic communities, and iii) identify ecological drivers of eukaryome structure and composition. Addressing these issues will help us to better understand the functional complexity of the coral holobiont and its possible responses to climate change.

## RESULTS

### Eukaryome diversity in coral reefs across the Pacific Ocean

To characterize the diversity, structure, and composition of the coral reef eukaryome, we collected 6,337 samples (coral, seawater, and sediment) spanning the tropical and subtropical reefs of the Pacific Ocean as part of the *Tara* Pacific expedition from 2016 and 2018 (Fig. 1A) (*37*, *40*). After quality control and filtering, we resolved 1.3 × 10⁹ 18S V9 sequences into 121,232 microeukaryote ASVs, including protists and small metazoans (Fig. 1, Supplementary Table 6). Over 80% of the ASVs detected in our survey (97,764 ASVs) were new compared to the *Tara* Oceans sampling (https://doi.org/10.5281/zenodo.18153978). However, the novel ASVs only represented 19% of the total number of sequences, indicating their overall low abundance.

**Figure 1.**
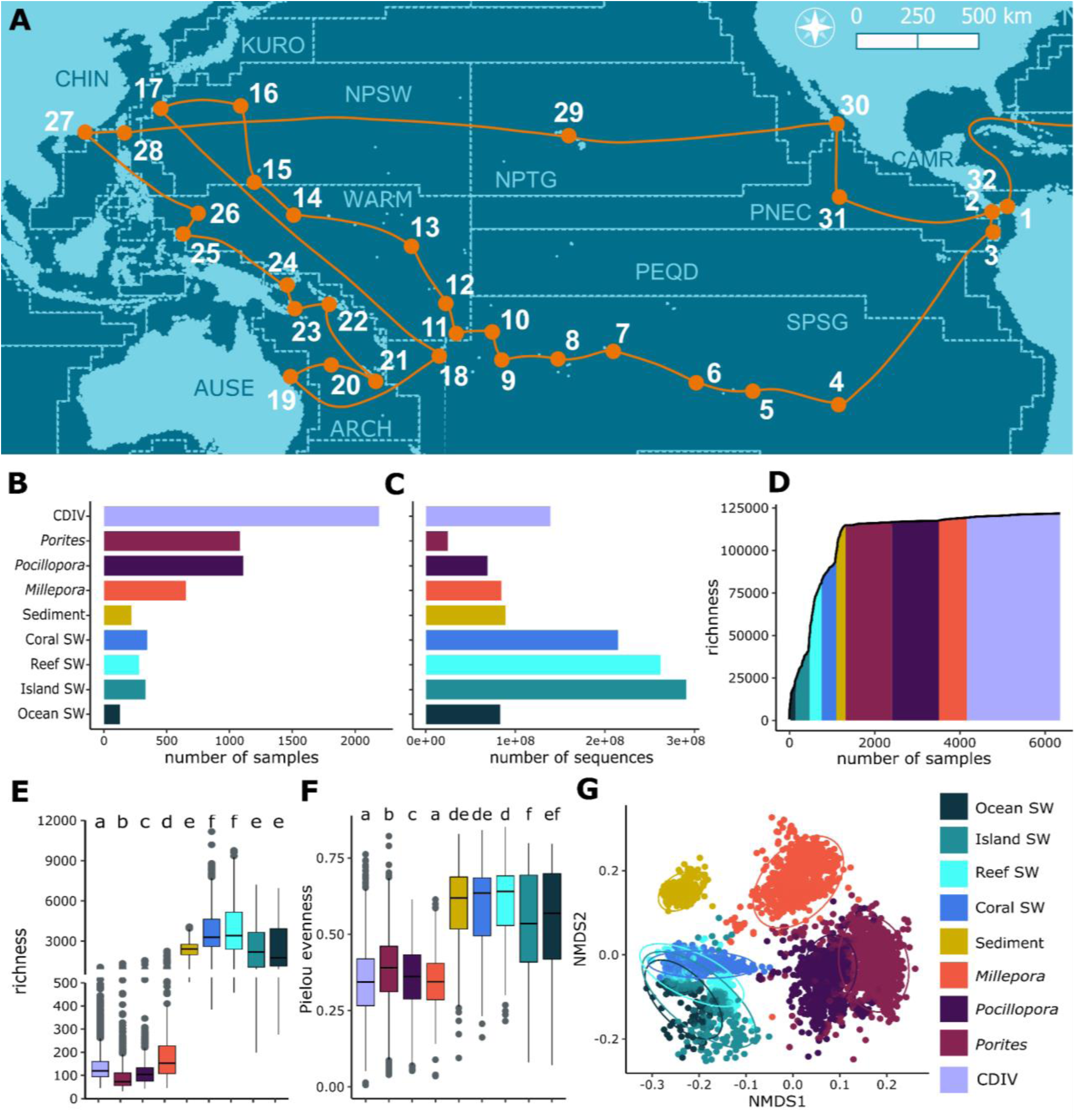
Overview of eukaryotic richness and diversity captured from stony corals, the surrounding seawater and sediment during the *Tara* Pacific expedition between 2016 and 2018. **(A)** Map of the *Tara* Pacific Expedition showing the island sites sampled in this study, numbered by order of sampling. Longhurst provinces are indicated by their code names. **(B)** Number of samples across sample types after quality filtering. (**C)** Number of sequences. (**D)** ASV accumulation curve in relation to sampling effort across sample types. **(E)** Observed richness across sample types. **(F)** Pielou’s evenness index across sample types. **(G)** nMDS ordination across sample types (excluding CDIV) based on Jaccard dissimilarity matrix (stress = 0.14). Colors are consistent across all panels. Samples in this study were taken in the Longhurst provinces of: Kuroshio current (KURO), China Sea (CHIN), Northwest Pacific subtropical (NPSW), Western Pacific warm pool (WARM), East Australian coast (AUSE), Archipelagic deep basins (ARCH), North Pacific Tropical gyre (NPTG), North Pacific equatorial counter current (PNEC), Pacific equatorial divergence (PEQD), South Pacific gyre (SPSG), and Central American coast (CAMR) (*41*).

Sampling across 113 coral reefs at 32 islands, yielded 5,035 coral samples (79.4 % of total), 1,083 seawater samples (17.1 %, comprising ocean, island, reef, and coral-surrounding SW based on their increasing proximity to the sampled corals), and 219 sediment samples (3.4 %). Seawater and sediment samples accounted for the majority of microeukaryotic diversity (92,747 and 49,246 ASVs, respectively), representing 92 % of all ASVs (Fig. 1D) and 74.9 % of the total number of sequences. Accordingly, richness in environmental samples ranged between 2,000-3,000 ASVs per sample, which was 20- to 40-fold higher than that of the coral samples (50-150 ASVs; Fig. 1E; pairwise p < 0.05; Supplementary Table 7A). Rarefaction curves (Supplementary Figure 1) indicate that seawater samples (except island seawater, SW), sediments, and pooled coral samples (CDIV) reached saturation. The total number of sequences in coral samples was low compared to the rest of the sample types (except ocean SW, Figure 1C), due to the co-amplification of 18S host sequences (Supplementary Table 2). Microeukaryotic communities of coral samples were also less even than those of the environmental samples (p < 0.05; Fig. 1F; Supplementary Table 5B), caused by the dominance of Symbiodiniaceae (75-89 % mean relative abundance) compared to seawater and sediment communities (28 % Symbiodiniaceae in coral SW, 0.03-0.3 % in the others). The overall most abundant ASV was also assigned to Symbiodiniaceae (3.6 % of total sequences), and the most prevalent ASV was assigned to the Symbiodiniaceae genus *Cladocopium* (89 % of all samples) (Supplementary Table 6).

Ordination analyses using non-metric multidimensional scaling revealed that the composition of the eukaryome differed between seawater, coral, and sediment samples, with little to no overlap between them (Fig. 1G, p < 0.05, Supplementary Table 8A, B). Water and sediment eukaryomes were more homogeneous than in coral samples (p < 0.05, Supplementary Table 8C). Within the seawater samples, the greatest overlap occurred among ocean, island, and reef SW, which separated into two groups according to size fraction (Supplementary Figure 2). Microeukaryote communities of coral surrounding seawater were more homogeneous and more similar to those of corals than to other environmental samples, forming a transition zone between the environment and corals, particularly *Pocillopora*, from which these samples were collected. Among coral samples, the microeukaryote communities of the scleractinian coral genera *Porites* and *Pocillopora* were more similar to each other than to the hydrozoan coral *Millepora* (Fig. 1G). Overall, the eukaryomes differed in diversity, evenness, and composition across coral, seawater, and sediment. Coral-associated communities were less rich and less even than those of seawater and sediment, and community composition also differed among coral genera.

### Coral-associated eukaryome

To further explore the coral-associated eukaryome across phylogenetically and functionally diverse host species, we collected two complementary sample sets. At three reefs per island, we systematically collected ten colonies of three coral genera (*Pocillopora*: 1,109 samples, *Porites:* 1,082 samples, *Millepora:* 652 samples). In addition, we sampled 80 coral colonies randomly to capture overall coral diversity at one reef per island (“CDIV”: 2,192 samples). These > 5,000 coral samples, representing 29 genera and taxon groups, were dominated by 25 eukaryotic lineages, which constituted over 75 % of the microbial communities, after removal of Symbiodiniaceae. The sponge class Demospongiae, the crustose coralline algae Corallinales, and Maxillopoda (primarily copepods and barnacles) were consistently among the most abundant taxa across all coral lineages. Other taxa showed strong host associations, including Annelida in *Cyphastrea*, *Diploastrea*, *Echinophyllia*, and *Leptastrea*; Gigartinales (Rhodophyta) in *Goniastrea*; Hildenbrandiales (crustose Rhodophyta) in *Seriatopora*; Homoscleromorpha (Sponge) in *Dispasastrea;* and unclassified eukaryotes in *Astreopora* and Dendrophyllidae (Fig. 2).

**Figure 2.**
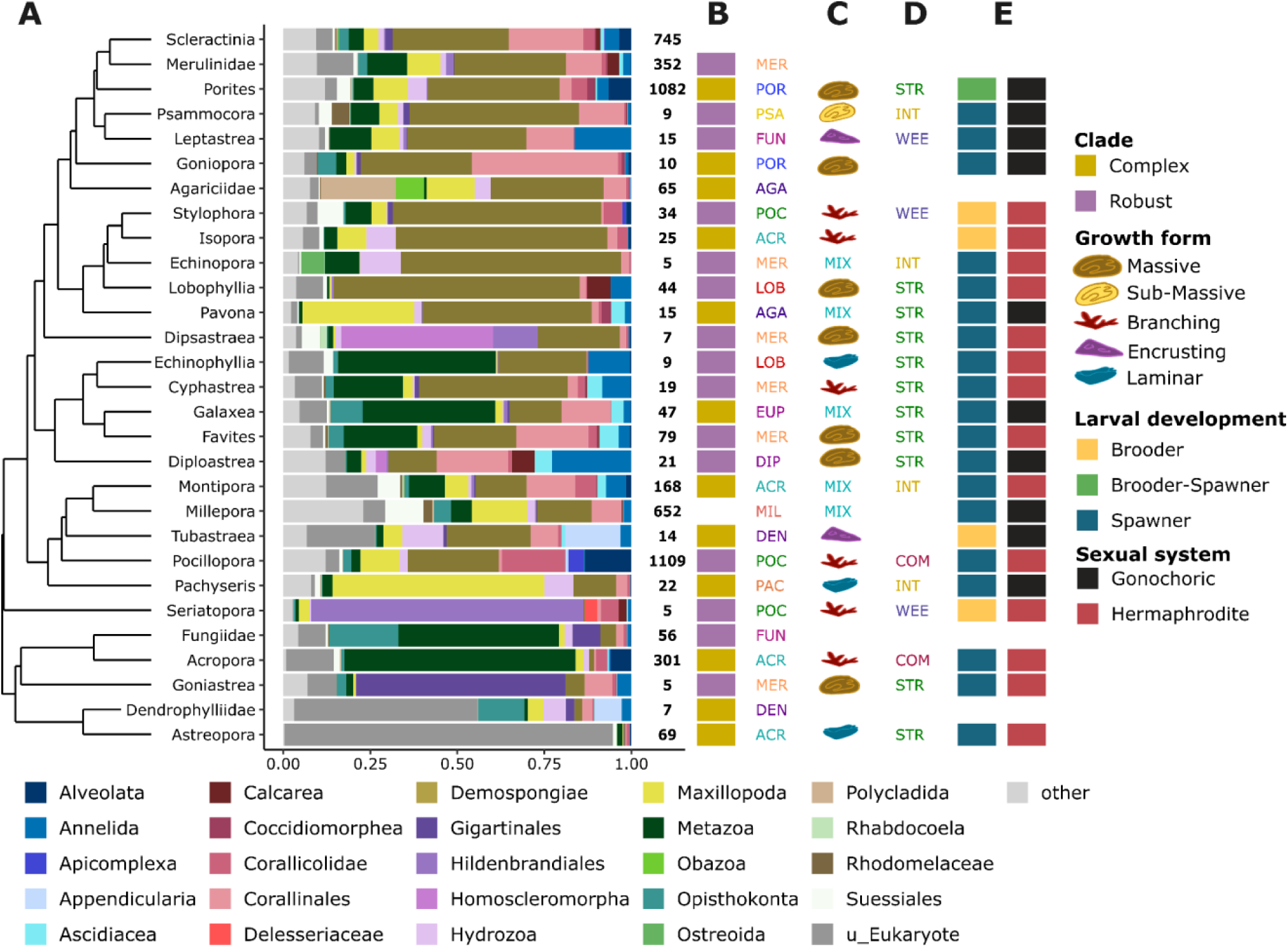
Microeukaryotic community, including small metazoans, and traits of sampled corals from the *Tara* Pacific Expedition (n ≥ 5). Corals were identified to the lowest possible taxonomic level (see Materials and Methods). **(A)** Corals were clustered by the Bray-Curtis similarity of their eukaryotic communities. Relative abundance of the lineages found in the corals sampled and the number of samples per coral. **(B)** Coral clades (Complex or Robust) and families, coded as the first three letters of the family name, are indicated. **(C)** Main growth form, being massive, submassive, branching, encrusting, laminar or a mix of several growth forms (MIX). **(D)** Life history traits of the coral taxa, as Stress-tolerant (STR), Competitive (COM), Weedy (WEE) or Intermediate (INT). **(E)** Larval development (Brooder, Spawner or mix), and sexual system (Gonochoric or Hermaphrodite).

To further explore the link between eukaryotic community composition and coral host traits, we clustered eukaryomes by Bray-Curtis similarity and juxtaposed the dendrogram with host traits, including coral phylogeny, taxonomy, growth form, life-history strategy, and reproductive mode. Eukaryomes of Pacific corals clustered into two major groups, which differed in the abundance of Demospongiae (Fig. 2A). In the smaller cluster, Demospongiae accounted for < 5 % of the microbial communities of the corals Fungiidae, *Acropora*, *Goniastrea*, Dendrophyllidae, *Astreopora,* and *Seriatopora*. The remaining 23 coral taxa harboured > 12 % relative abundance of Demospongiae. Despite this partitioning, overall community similarity showed little correspondence with host phylogeny or functional traits, including coral taxonomy at the clade or family level (Fig. 2B), growth form (Fig. 2C), life history traits (Fig. 2D), or reproductive mode (Fig. 2E). The microeukaryotic community composition of *Millepora*, the only hydrozoan, was similar to that of the scleractinian corals and clustered with *Montipora*. Overall, coral-associated eukaryotic communities were dominated by a few consistent taxa, particularly Demospongiae, Corallinales, and Maxillopoda. This pervasive dominance of small metazoans was unexpected and contrasts with the protist-centered view of coral-associated microeukaryotes. Community composition did not show clear links with coral host traits, but differed notably in the relative abundance of Demospongiae. Because of uneven sampling of most coral lineages across the expedition, all subsequent statistical analyses focused on the systematically sampled coral genera *Millepora, Pocillopora*, and *Porites*.

### Coral reef habitats as hotspots of microeukaryote diversity

Next, we aimed to assess distribution and partitioning of the microeukaryotic communities across seawater, sediment, and host-associated habitats in Pacific coral reefs. Across the 52 microeukaryotic lineages, many were overrepresented in or even exclusive to one habitat. Moreover, the correlation between ASV abundance and diversity was moderate (R²adj = 0.49; Supplementary Figure 3), and many lineages were more abundant in one habitat, but more diverse in another. Seawater harboured several phylogenetic lineages that were rare or absent in other compartments (e.g., Radiolaria and the flagellates Parabasalia; Fig. 3). Sediments contained both unique lineages (e.g., Rotosphaerida, Preaxostyla) and disproportionately high diversity despite low abundant lineages such as Bigyra, Haptista, Kathablepharida, Cryptophyta, and some Excavata/Metamonada. Corals overall harboured fewer and less diverse lineages than environmental samples, consistent with their lower richness (Fig. 1D), except for the Pluriformea, for which *Millepora* hosted all the diversity of the lineage (4 ASVs). Two lineages, Dinoflagellata (16,503 ASVs), including Symbiodiniaceae, and Metazoa (22,327 ASVs), stood out with exceptionally high diversity. The prominence of metazoan diversity across all reef compartments was particularly notable, together with recurrent but less diverse lineages, including the heterotrophic protists Apusomonadida (Obazoa) and Centroplasthelida (Haptista), these eukaryotic lineages were consistently present across most reef habitats. Unclassified Eukaryotes (u_Eukaryote) represented ∼25 % of ASVs and were widely distributed, particularly abundant, and diverse in seawater and sediment samples, suggesting that a substantial fraction of reef-associated eukaryotic diversity remains evolutionarily unresolved.

**Figure 3.**
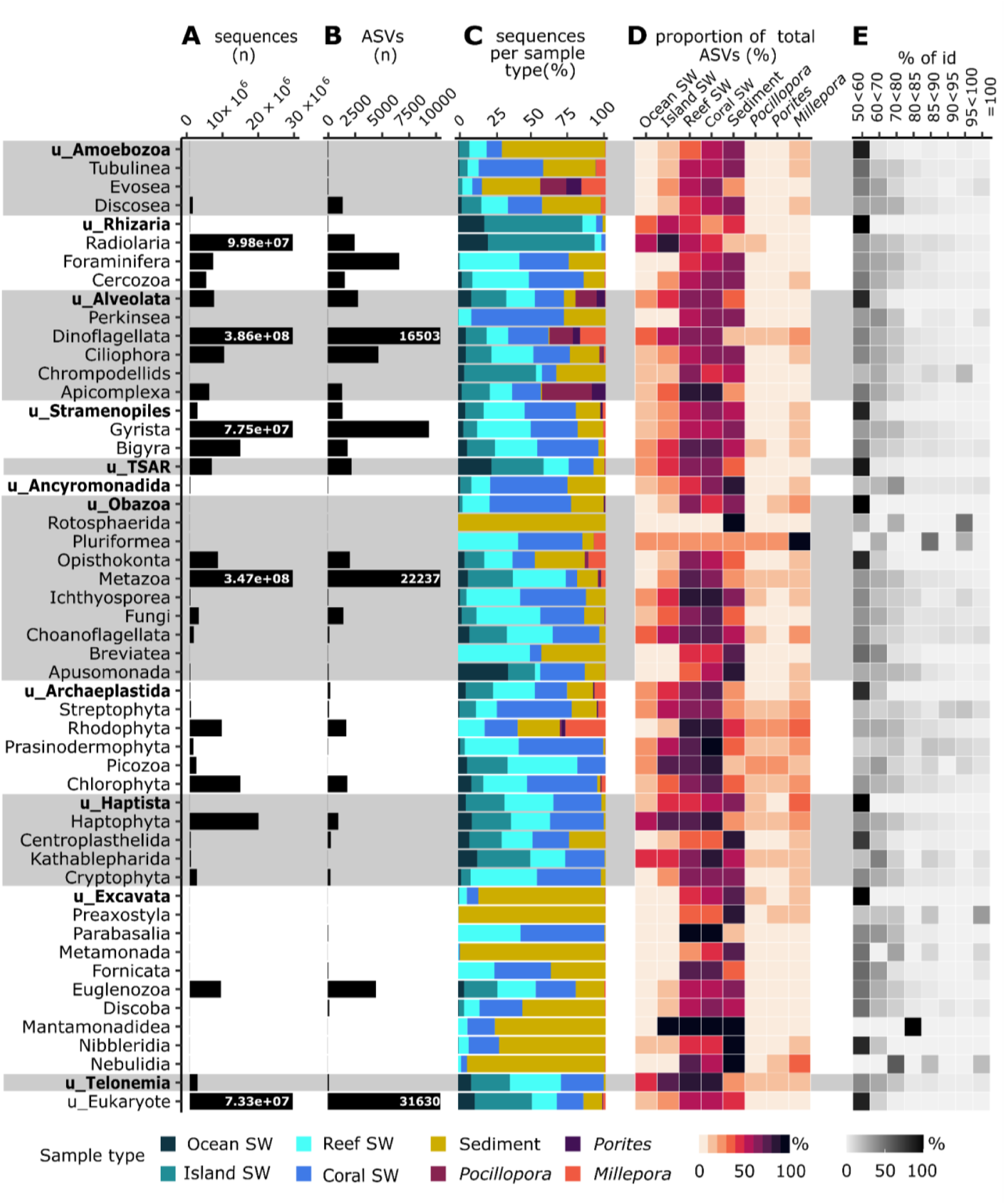
Overview of the main eukaryotic lineages, including small metazoans, observed across sample types: ocean, island, reef and coral seawater, sediments, and the coral *Millepora*, *Pocillopora*, and *Porites* from the *Tara* Pacific expedition. In bold is the name of the major lineage, and it includes all unclassifiable ASVs for that group. Below the major lineages are their sub-lineages. **(A)** Number of sequences per lineage, bars are numbered when they exceed the axis. **(B)** Observed richness as the number of ASV per lineage, bars are numbered when they exceed the axis. **(C)** The proportion of sequences for each lineage found in each sample type. **(D)** The percentage of distinct ASVs from each lineage that are present in each sample type. **(E)** Percentage of ASVs assigned to the closest reference based on similarity between 50–60 %, 60–70 %, 70–80 %, 80–85 %, 85–90 %, 90–95 %, 95–100 %, and 100 %.

This study revealed a high proportion of cryptic and novel diversity. Most of the ASVs were assigned to a low number of reference sequences (Supplementary Figure 4) with low similarity (50-70 %; Fig. 3E), and only 2,277 ASVs assigned with more than 95 % confidence. This pervasive lack of taxonomic resolution highlights the major underrepresentation of reef-associated microeukaryotes in current databases, although it may also partly reflect the limited phylogenetic resolution of the 18S V9 marker (*42*). Most eukaryotic lineages were detected in almost all Longhurst provinces, but those that were found in fewer provinces were always associated with sediments (e.g., Rotosphaerida, Praxostyla, Metamonada) (Supplementary Figure 4C). These results show that microeukaryotic lineages have distinct patterns of abundance and diversity across seawater, sediment, and coral habitats. A large proportion of the detected ASVs showed low similarity to reference sequences, highlighting the extent of uncharacterised eukaryotic diversity in coral reef ecosystems.

### Biogeographic patterns in microeukaryotic community composition

To examine how the composition of the coral reef eukaryome varied across the Pacific Ocean and whether geographically closer islands hosted more similar communities, we compared the eukaryome composition across sampling locations. Consistent with divergence of corals across the Pacific Ocean (*43*), we observed biogeographic partitioning of coral-associated microeukaryotes. Eukaryomes in the Tropical Eastern Pacific (islands 1-3, 31-32) were composed of distinct communities compared to the central, southern, and western regions (Fig. 4, Supplementary Figure 5). However, within and across all sample types and this regional structuring, geographic proximity between islands did not predict community similarity, as indicated by the distance decay analysis (Mantel ρ ranging from -0.361 to -0.037, all p > 0.05; Supplementary Figure 6). Across the Pacific, the eukaryome of the three corals shifted towards a higher proportion of Maxillopoda in the western and southwestern Pacific provinces (islands 8 to 28).

**Figure 4.**
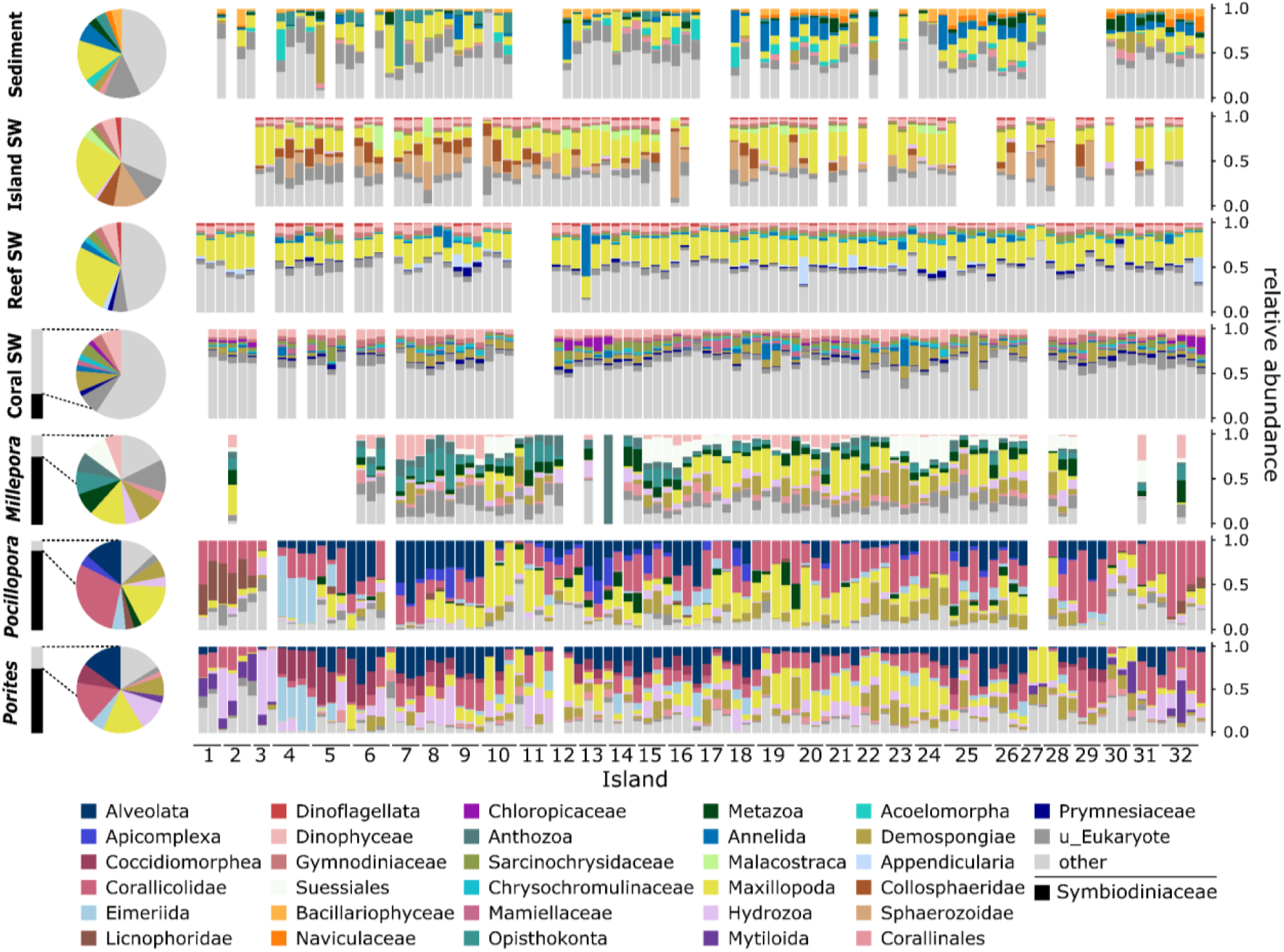
Relative abundance of microeukaryotic lineages, including small metazoans, in the different sample types across island sites of the *Tara* Pacific expedition. To improve visualisation in corals and surrounding seawater (Coral SW), Symbiodiniaceae ASV abundances are plotted separately, as they represent a large proportion of sequences (left). Pie charts summarise the mean relative abundance of each lineage across all sites for each sample type. Bar plots show the mean relative abundance per island site, where each bar represents one sampling site. Island numbers correspond to the sampling map in Figure 1A. Sites where only corals or only the environment were collected were excluded (102/113 sites kept). Replication per site, coral species n = 10, sediments n = 3, coral SW n = 4, reef SW n = 3, island SW n = 4.

Overall, microeukaryotic communities of seawater and sediment were more consistent between sampling stations than the variable communities captured in the corals. Maxillopoda maintained a stable, high relative abundance in seawater and sediment samples across the entire Pacific (13-26 % mean abundance), except for the coral SW, in which Maxillopoda had low abundance (<1 %). Reef SW was additionally dominated by Dinophyceae, the brown algae Sarcinochrysidaceae, and unclassified Eukaryotes (6-3 % mean abundance), while island SW contained higher proportions of radiolarian lineages (Collosphaeridae and Sphaerozoidae), but only in the temperate open ocean stations (islands 4-11 and 18). Sediments contained higher proportions of diatoms (Bacillariophyceae and Navicula) and Annelids, and included Demospongiae, which were also present in coral SW and corals (Fig. 4). Together, these results indicate that habitat type exerted a stronger influence on microeukaryotic community composition than geographic distance. Coral-associated eukaryomes were markedly more variable across the Pacific than seawater or sediment communities, suggesting weaker environmental homogenisation and stronger host- or locality-specific assembly processes.

### Habitat specificity and ubiquity of the eukaryome

To further assess habitat specificity of the eukaryomes, we examined the overlap of ASVs across sample types and identified the most prevalent taxa within each coral genus. Eukaryotic communities were highly habitat-specific with one-third of ASVs occurring in only one sample type (38,239 ASVs) (Fig. 5A). Of these, more than half (20,561 ASVs) were unique to sediments, representing 40 % of the ASVs in this habitat. The open ocean system (island and ocean SW) shared 4,863 exclusive ASVs, while coral reef samples (reef SW, coral SW, sediments, and corals) together shared 112,493 ASVs, which accounts for 94 % of the total diversity found. In contrast, only a small number of ASVs (n = 1,422) were shared across all sample types, and < 1,300 ASVs were exclusively shared among the three corals, representing only 5 % of their combined ASV diversity. This limited overlap indicates strong ecological partitioning of reef-associated microeukaryotes across environmental and host-associated compartments.

**Figure 5.**
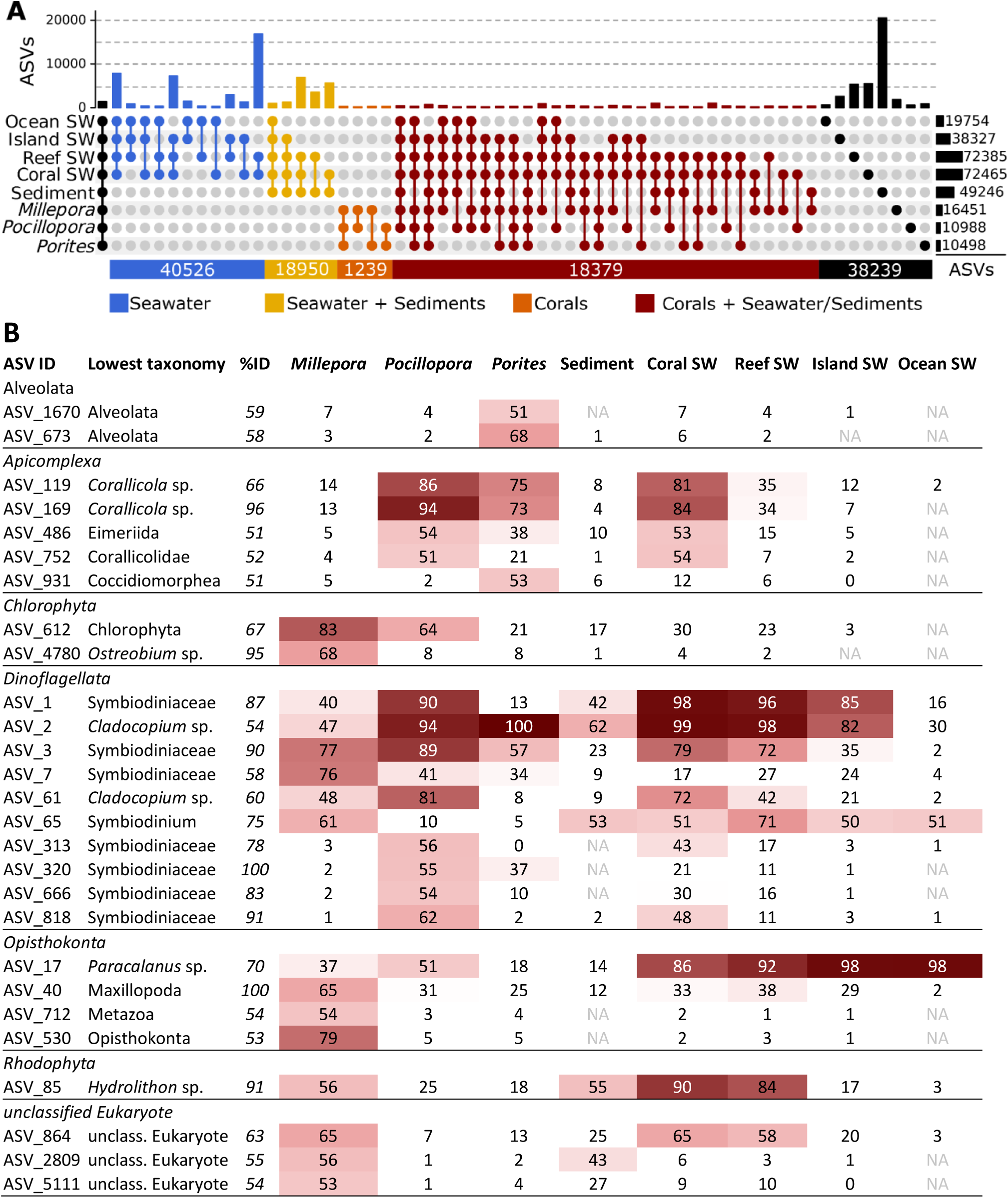
Shared and most prevalent ASVs found in stony corals and their surrounding seawater and sediments during the *Tara* Pacific expedition. **(A)** Upset plot of the shared ASVs between sample types (method distinct). Only intersections above 200 ASVs have been plotted. Intersections that only include seawater are in blue, only include sediments and seawater in yellow, only corals in orange, and corals with sediments and/or seawater in red. **(B)** Prevalence of ASVs present in more than 50% of *Millepora*, *Pocillopora*, or *Porites* samples, across all sample types. ASVs that were not present in a sample type are represented with *NA*. The ASV name is correlated with their abundance in the whole dataset (e.g. ASV_2 is the second most abundant ASV in the dataset). ASVs are grouped by taxonomic ranks, and the %ID column indicates the confidence of that taxonomic assignment to the reference database.

Host filtering was also apparent in prevalence patterns, in which the most prevalent ASVs (present in > 50 % of coral samples) were generally less prevalent or absent in the surrounding environment (Fig. 5B), except for the coral SW (e.g., Apicomplexa, *Paracalanus*, or Rhodophyta). Apicomplexans specifically, were only prevalent in scleractinian corals and coral SW (Fig. 5B). *Pocillopora* had the highest number of prevalent ASVs (14 ASVs), with *Corallicola* (ASV_169) and *Cladocopium* (ASV_2) present in 94 % of samples. *Millepora* had 13 prevalent ASVs, many of which were also common in seawater or sediment samples (e.g., ASV_85, *Hydrolithon*). *Porites* had seven ASVs present in > 50 % of samples, with *Cladocopium* (ASV_2) occurring in 100 % of samples. In summary, microeukaryotic communities were strongly habitat specific, with very few ASVs shared among sample types. Within corals, a small core assemblage of prevalent ASVs, dominated by Symbiodinicaeae and Apicomplexa, persisted across colonies, supporting stable and functionally important host-associations.

### Network of the Pacific eukaryome

To explore connections between the occurrence of microeukaryotic ASVs and the structure of the Pacific coral reef eukaryome, we constructed co-occurrence networks for the full dataset and separately for each coral genus. The Pacific coral reef Eukaryome formed a large, but weakly interconnected network structured by habitat type. The network was composed of Alveolata, Dinophyceae, the radiolarians Sphaerozoidae, Opisthokonta, unclassified Metazoans, Demospongiae, and Maxillopoda (mainly copepods and barnacles), representing 18 % of all ASVs in the dataset (21,862 ASVs) (Fig. 6). In line with the presence of subcommunities associated with specific sample types and an intermediate modularity (0.59), the network was overall highly disconnected (edge density: 0.0003). Island seawater formed a more distinct and disconnected network dominated by Sphaerozoidae. Sediment clusters were closer to coral surrounding seawater than to the rest of the sample types. Among coral genera, *Pocillopora* and *Porites* networks were closely connected (Fig. 6, Supplementary Figure 7).

**Figure 6.**
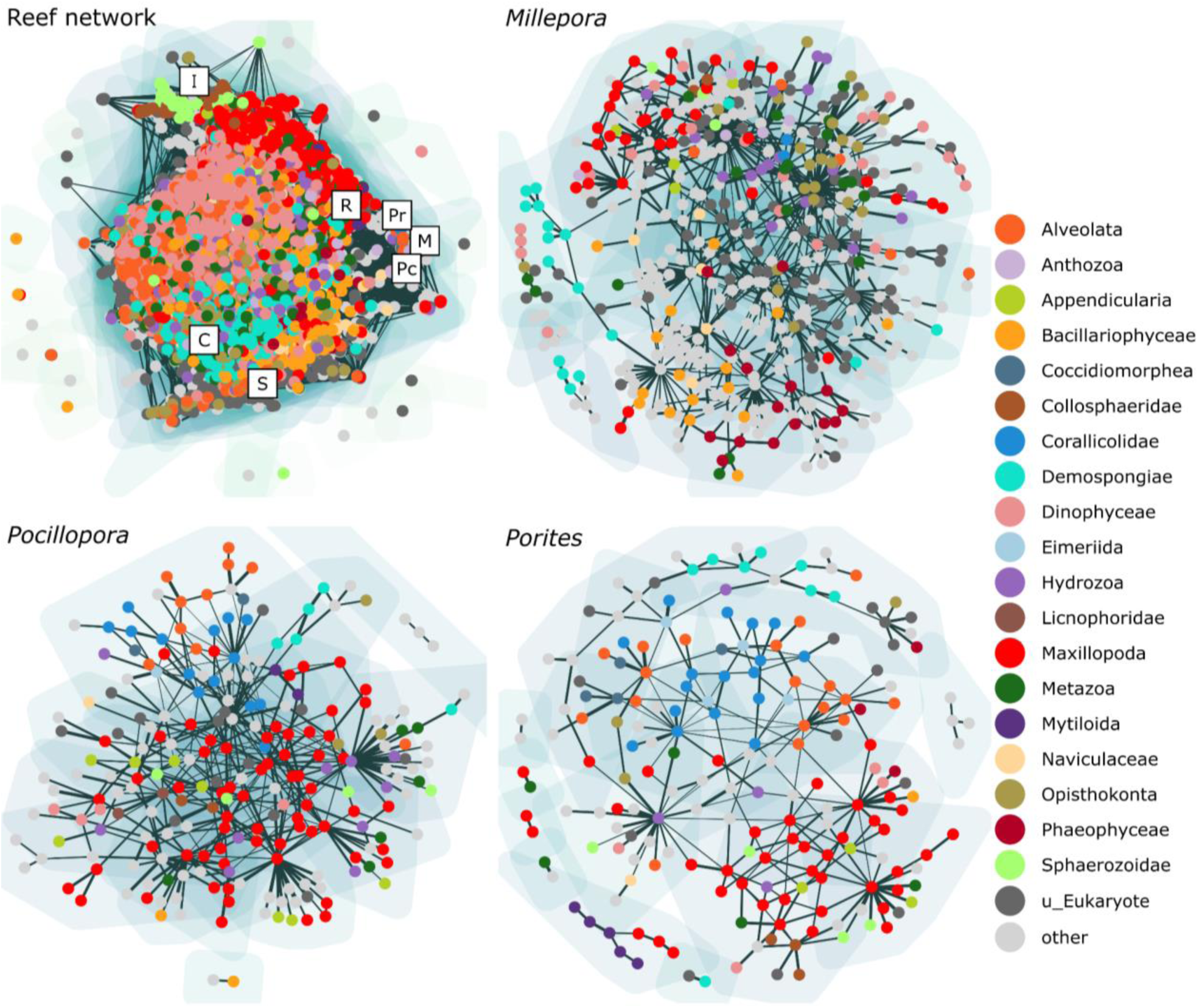
Network of the reef eukaryome (seawater, sediments and coral samples) and within the corals sampled, with each sample type labelled in their centroid - *Millepora* (M), *Pocillopora* (Pc), and *Porites* (Pr), ocean (O), island (I), reef (R) and coral surrounding (C) seawater, and sediments (S). The coral networks do not include Symbiodiniaceae. Nodes represent unique ASVs and are colored by the lowest taxonomic rank identified. Lines connect nodes with positive associations, and line width is related to the strength of that relationship. Shades represent clusters of ASVs, calculated with Louvain.

When analysed separately, each coral showed a distinct and compartmentalised network structure (modularity: 0.65-0.77). Corresponding to the differences in the total number of ASVs found per coral, *Millepora* had the largest network, followed by *Pocillopora* and *Porites* (461, 228, and 191 ASVs, respectively). Their eukaryome networks had low overall connectivity (edge density: 0.006, 0.015, and 0.013 for *Millepora, Pocillopora,* and *Porites*, respectively). *Pocillopora* had the highest number of connections per ASV, followed by *Millepora* and *Porites* (mean degree: 3.4, 3.0, and 2.6, respectively), although this was lower than the average number of connections in the Pacific network (5.79).

Within coral eukaryomes, associations occurred mainly between ASVs from different taxonomic lineages, forming highly mixed groups (assortativity: 0.164, 0.259, and 0.290 for *Millepora*, *Pocillopora*, and *Porites*, respectively). In addition, highly connected ASVs tended to link less connected ASVs, acting as key ASVs in the community structure (assortativity degree: -0.212, -0.294, and -0.294 for *Millepora*, *Pocillopora*, and *Porites* respectively). These key ASVs mainly belonged to Maxillopoda and the apicomplexan Corallicolida in *Pocillopora* and *Porites*, while in *Millepora*, they belonged to Maxillopoda, the diatom Naviculae, and other taxa (Fig. 6; Supplementary Figure 8). Despite their abundance and prevalence in the coral eukaryome, Demospongiae were only found at the periphery of the coral networks. Together, these results show that the Pacific coral reef eukaryome forms a large but weakly connected network structured by sample type. Within corals, network organisation was dominated by a limited number of recurrent and highly connected taxa, particularly Maxillopoda and Corallicolida, highlighting their potential ecological importance in coral-associated microeukaryotic assemblages.

### Environmental drivers of the Pacific reef eukaryome

To identify the environmental factors driving the eukaryome composition, we used distance-based redundancy analysis with a set of 45 environmental composite variables including temperature, salinity, seawater chemistry and historical stress indices among others. Environmental variables explained between 13-48 % of variation in eukaryotic community composition across sample types. The highest proportion of variance explained by environmental variables was observed in *Millepora* (48 %) and sediments (40 %), followed by *Pocillopora* (34 %), coral SW (28 %), *Porites* (25%) and reef SW (21 %). For the island (13 %) and ocean (21 %) SW eukaryomes, less variance was explained by the environment, with the lower values likely due to a lower number of available explanatory variables such as the absence of long-term SST variables.

Across corals, temperature-associated variables were consistently the strongest predictors of community composition, although their relative importance varied among hosts (Fig. 7, variable names in Supplementary Table 4 and 5). Sea surface temperature (SST) was one of the major drivers across the three corals, being most important in *Pocillopora* (42 %) and ranked second in *Millepora* (21 %) and *Porites* (18 %). Cold stress metrics showed the highest correlation in *Millepora* (62 % for degree cooling week frequency). Eukaryotic communities in *Pocillopora* and *Porites* also correlated with cold variables, though less strongly (e.g., 28 % for average cold Thermal Stress Anomaly frequency in *Pocillopora*, and 16 % for Thermal Stress Anomaly degree cooling week frequency in both). Sea surface temperature anomalies also had a high correlation in *Pocillopora* (26 %). Furthermore, sea surface salinity (SSS) had a consistent correlation with eukaryomes of *Millepora* (35 %) and *Pocillopora* (23 %) (Fig. 7). Spatial variables such as latitude and longitude were significant only in *Millepora* (23 % and 14 %), which is in line with its slightly higher distance-decay than in the other corals (Supplementary Figure 6). Finally, from all chemical parameters included, only pH and silicate concentration were moderately correlated with community composition in *Millepora* and *Pocillopora* (18–11 %), but not in *Porites* (5 %).

**Figure 7.**
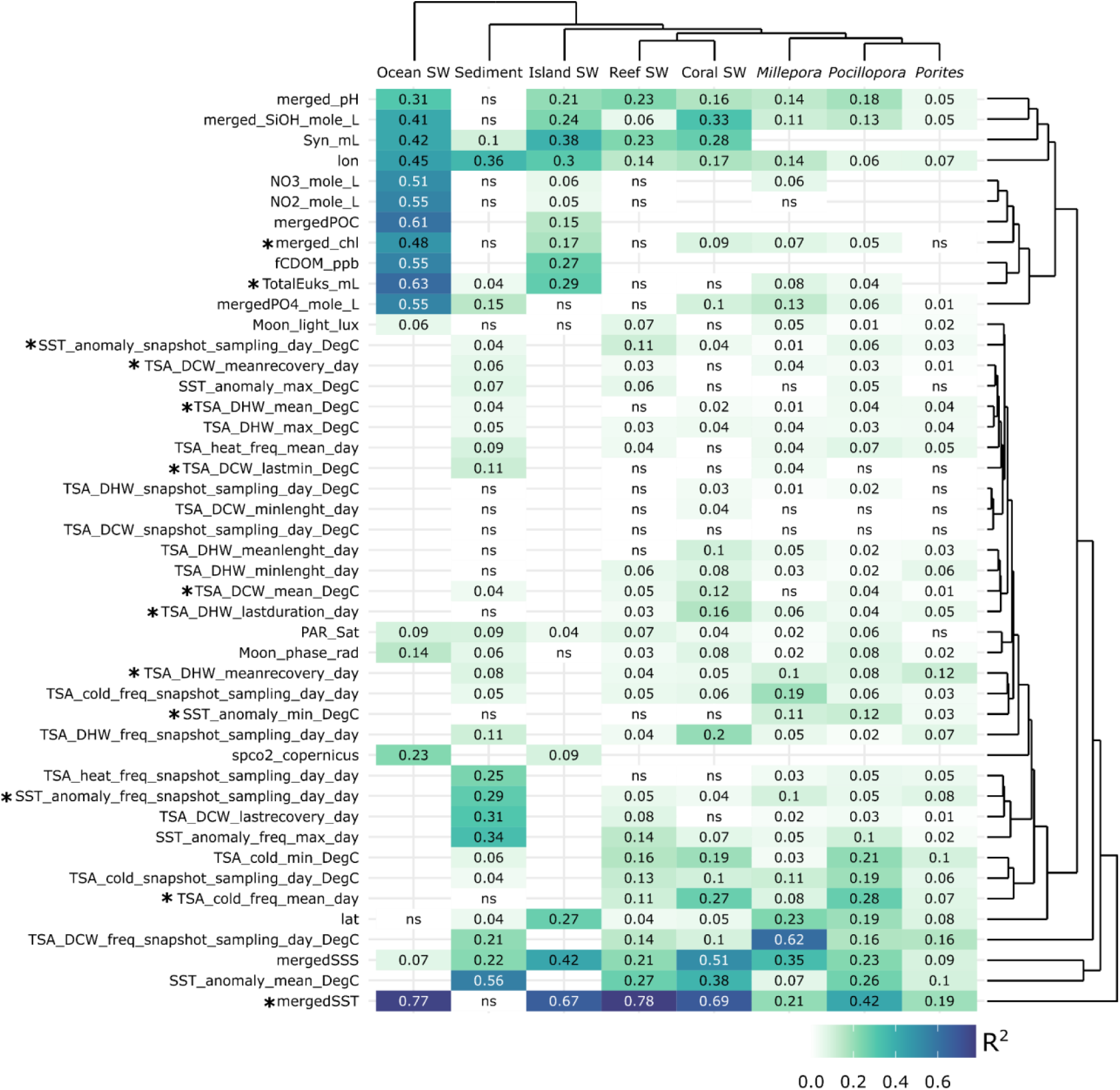
Correlation between the environmental variables and the community composition of the different sample types of the *Tara* Pacific expedition. Colours correspond to the value of R2. Only significant variables have the R2 value indicated, and only variables with 90% or more of the values were included per sample type. Correlations were calculated by fitting the environmental variables to the distance-based RDA matrix (Supplementary Figure 9). Hierarchical clustering of variables and hosts was done with *UPGMAM.* Full names, descriptions, and source datasets of the environmental variables can be found in Supplementary Table 4. For correlated variables, one representative variable was selected (∗); the full list of correlated variables is provided in Supplementary Table 5.

In the seawater, planktonic communities were strongly structured by sea surface temperature and salinity, together with chemical parameters (e.g., pH, and silicate, nitrate, and nitrite concentrations) (Fig. 7). These correlations were typically strongest in the open ocean, slightly weaker in island SW, and became much less relevant in reef SW, coral SW, and sediments. For instance, pH ranged from 31 % correlation in ocean SW to non-significant in sediments, nitrate ranged from 51 % in ocean SW to 6 % in island SW and non-significant in reef SW and sediments (Fig. 7). Overall, these results show that temperature was the dominant driver of coral-associated microeukaryotic community composition, with salinity and chemical parameters playing secondary roles. The strong influence of thermal stress variables across coral hosts further suggests that reef eukaryomes are highly sensitive to ongoing climate-driven environmental change.

## DISCUSSION

This study provides a Pacific-wide baseline for coral- and reef-associated eukaryomes. Coral-associated communities were host-specific, with unique community compositions and a large number of unique ASVs, suggesting that the loss of coral hosts could lead to the disappearance of their associated eukaryotic communities. These coral-associated communities were diverse and uneven, composed of co-occurring and highly prevalent ASVs, and dominated by a few abundant lineages, including microeukaryotes such as Apicomplexa, as well as previously overlooked metazoans belonging to Maxillopoda and Demospongiae. Environmental factors, in particular SST and the frequency of cold stress events, were more strongly correlated with community composition than heat anomalies, bringing attention to marine cold spells in coral reef ecology. Finally, we uncovered an interconnected Pacific-wide community, in which different habitats fill complementary roles for specific lineages, some act as diversity reservoirs, harbouring many distinct ASVs of a lineage, while others act as biomass reservoirs, concentrating most of its abundance. Together, these findings establish a baseline for understanding reef microeukaryotic and small metazoan diversity and highlight the vulnerability of the coral eukaryome in a rapidly changing ocean.

### Unprecedented eukaryome diversity of Pacific reefs

We recovered 121,232 eukaryotic ASVs from the corals, seawater, and sediments, an exceptionally high and previously undocumented diversity of reef-associated microeukaryotes, and highlighting coral reefs as hotspots of eukaryome diversity. The number of eukaryotic taxa we found in this unprecedented survey effort of over 6,300 samples across Pacific coral reefs, is almost triple the diversity found in other global coral surveys (*32*). Compared to the *Tara* Oceans sampling we detected 1.5 times the global planktonic diversity (of 335 samples) when comparing OTUs (226,759 OTUs here vs. 150,000 OTUs in de Vargas et al. 2015), 80 % of which was previously unrecorded (*44*). Based on similar sampling efforts, ASV richness in our reef sediment samples alone (219 samples) was eight times that of global soil eukaryotes (*45*).

To date, the microeukaryotic communities of only a few biomes have been comprehensively studied, and our results highlight how much more of the eukaryome diversity remains to be discovered. Even this extensive survey likely captures only a fraction of total reef-associated diversity, with potentially more diversity to discover in this biome, as suggested by the rarefaction curves. Dinoflagellates stood out as exceptionally diverse but remain underrepresented in reference databases (PR2 2 v.5.0.0), highlighting that the taxonomic assignments in this study should be interpreted as a broad overview rather than precise lineage-level identification. As we continue to revise global estimates of microbial diversity (*9*, *46*, *47*) future work will expand our understanding of these communities and their ecological roles by implementing host-excluding primers (*14*, *48*) and lineage-specific markers (*49*) as well as databases to improve taxonomic resolution.

### Host-specific and uneven microeukaryotic communities in Pacific corals

Corals not only are the cornerstone of highly diverse reef ecosystems, but they also provide a habitat for a rich community of eukaryotic organisms beyond Symbiodiniaceae (*50*). Corals harboured distinct eukaryomes, yet these differences were not explained by coral taxonomy, life traits, or morphology, which contrasts to the patterns driving coral-associated prokaryotic communities (18, 51, 52). This weak correspondence with host phylogeny suggests that coral-associated eukaryomes may be shaped by finer-scale ecological or physiological filters acting at the species or colony level. Similar, non-phylogenetically correlated eukaryomes have been reported in other phyla, from marine sponges to non-human primates (*53*, *54*).

Despite differences in the community composition, corals had similar eukaryome structure dominated by a few highly abundant taxa, a trend also seen in coral-associated prokaryotes (*9*, *19*, *55*). The consistent association with Demospongiae, Corallinales, Maxillopoda, and other Metazoans across all corals points to non-random relationships with the host encompassing parasitism, mutualism, inquilinism, and predator-prey dynamics (56–58). The widespread dominance of metazoan lineages was particularly unexpected and substantially broadens the current view of coral-associated microeukaryotic assemblages, which has largely focused on protists. However, clarifying these diverse roles will require functional assays and targeted genomic analyses (*31*). In addition, some of the most prevalent taxa identified in this study, including copepods, dinoflagellates, and ciliates, are known to have exceptionally high 18S rRNA gene copy numbers, which can inflate their apparent abundance in metabarcoding datasets (59, 60). Consequently, relative sequence abundance should not be interpreted directly as organismal biomass or ecological dominance. Functional and quantitative approaches will be needed to confirm the ecological roles inferred here.

The consistent Pacific-wide sampling of *Millepora, Pocillopora,* and *Porites* provides the first basin-scale catalogue of their eukaryomes. These three corals harboured distinct communities, with substantial variation, that did not follow a geographical signal. The hydrozoan *Millepora* hosted the most diverse eukaryome and acted as a hotspot for rare microeukaryotic lineages, including Pluriformea (*61*). In contrast, the scleractinian corals *Pocillopora* and *Porites* had less diverse and more even communities, and were less variable than *Millepora*, even though the latter consisted of fewer host lineages (*43*). These differences might be related to their distinct evolutionary histories, colony growth form, skeletal structures, and intrinsic susceptibility to colonisation (*62–64*). As expected only a small fraction of total ASVs were retained in the co-occurrence networks. This pattern indicates that coral eukaryomes combine a relatively small, consistently detected core, potentially enriched in symbiotic taxa, alongside a more variable component that may shift with environmental or host physiological conditions (e.g. (*65*).

Notably, previously identified coral-associated Apicomplexans such as Corallicolida were highly prevalent in *Pocillopora* and *Porites* (*30, 31*). Corallicolida have recently gained attention due to their widespread distribution across healthy and diseased scleractinians, though their ecological roles remain to be deciphered (*27*, *66*, *67*). Other putative health-related taxa were also observed, including the ciliate Licnophora in *Pocillopora* (*25*) and the bivalve Mytiloida in *Porites,* which have been shown to be associated with diseased corals and to debilitate coral skeletons (*68*). Interestingly, these were groups more abundant in the Central American region (CAMR), potentially indicating local stressors (*69*, *70*).

### Habitat partitioning of microeukaryotic communities

Microeukaryotic communities differed in composition and structure across habitats. Seawater communities and sediments had the highest richness and evenness, in contrast to less diverse and uneven coral communities. In addition to previously reported strong dispersal limitations in open-ocean microeukaryotic communities (*71*), the high number of unique ASVs detected in each habitat indicates strong habitat partitioning and niche specialisation in coral reefs, even within the same reef and geographic area. Such habitat specificity suggests that reef microeukaryotic diversity is maintained by fine-scale environmental filtering and ecological specialisation across reef compartments.

Microeukaryotic plankton communities were shaped by their distance to the reef system. Open ocean and island seawater were similar and both dominated by previously highlighted planktonic lineages such as Radiolaria (de Vargas et al. 2015). A clear community shift was observed towards reef seawater, where coral-associated groups, including Apicomplexa, Demospongiae, and Hydrozoa, were more prevalent. The coral surrounding seawater represents a microbial transition zone between the reef water and the coral host (*72*, *73*). Similar to the coral mucus layer, which it borders, it also reflects the composition of the underlying tissue-associated microbiome (*74*, *75*). Interestingly, the coral surrounding seawater was the only habitat lacking the otherwise abundant Maxillopoda, potentially reflecting their life history including a free-living larva that directly transitions to the coral tissue and establishes within the colonies (*76*).

Sediments acted as reservoirs of diversity and abundance of several microeukaryotic lineages, particularly members of Excavata, Centroplasthelida, Pluriformea, and Rotosphaerida. The ecological roles of these groups remain unclear but may involve participation in the microbial loop and nutrient cycling, as sediments are key in remineralisation processes (*77*). A notable fraction of sequences from sediments had no close match in reference databases, suggesting that reef sediments may harbour undescribed or poorly represented lineages (*78*). However, sediments also accumulate detrital and decaying material over long periods of time, which makes them biodiversity archives but might lead to an overestimation of present biodiversity (*79*).

### Environmental drivers and vulnerability of the eukaryome

Environmental variables strongly correlated with eukaryotic community composition. SST emerged as the most consistent predictor across habitats, except for sediments. SST is a major structuring force in marine ecosystems (*80*), acting as a global driver of surface ocean prokaryotic communities (Sunagawa et al. 2015), as well as both directly on organismal physiology and indirectly through effects on oxygen solubility and nutrient cycling (*81*, *82*). Despite the broad geographic scale, eukaryotic community composition showed only weak spatial structuring, which is in contrast to the corresponding prokaryotic microbiome recorded from the same samples (*9*). Instead, coral eukaryomes appeared more tightly linked to local environmental histories, particularly thermal regimes including cold anomalies, which were strongly correlated with *Millepora* and *Pocillopora* eukaryomes. This suggests a strong sensitivity, not only to heating, but also to cooling events and highlights the need to consider marine cold spells (*83–85*) because their frequency is projected to change under climate change with unknown implications for coral reef stability (*83–85*).

The correlation of SSS with the eukaryotic community composition of *Pocillopora* and *Millepora* corals agrees with large-scale analyses of oceanic plankton communities (*86*). Salinity fluctuations can also affect the host, as they can constrain coral thermal tolerance and survival (*87–89*). In contrast, seawater chemistry (e.g. silicates, nitrogen species, pH) had stronger correlations with planktonic than with coral-associated communities, likely reflecting the regulatory capacity of the host (*90*, *91*).

Network analysis revealed strong host structuring within the coral-associated eukaryome. The microeukaryotic network of *Pocillopora* was most connected and least modular, suggesting a more cohesive and potentially resilient community—as suggested previously for the *Pocillopora* microbiome (*92*, *93*)—compared with the more fragmented networks of *Porites* and *Millepora* (*94*, *95*). The conserved Pacific microeukaryotic network suggests the existence of a Pacific reef-scale eukaryome that extends beyond the previously described open ocean network (*86*). The presence of interconnected reef- and basin-scale networks suggests the existence of reservoirs for some of the taxa to recolonize adjacent habitats after disturbance. Highly connected keystone taxa likely contribute disproportionately to the stability and resilience of these coral–microeukaryote networks (*96*, *97*).

Coral reefs, which support nearly one billion people and a quarter of all marine life, have now surpassed a critical tipping point (*5*). As one of the most biodiverse ecosystems on Earth, understanding coral reef complexity is essential for mitigating its ongoing decline as well as the consequences thereof. Corals not only shape highly diverse reef environments, but also harbour within themselves a diverse community of organisms, including both prokaryotes and eukaryotes. By revealing the extent, structure, and environmental sensitivity of this hidden eukaryotic diversity, our study provides a foundation for integrating microbial eukaryotes into future models of coral reef resilience and ecosystem change.

## METHODS

### Sampling

The *Tara* Pacific Expedition (2016–2018) collected coral, seawater, and sediment samples at 113 coral reefs across 32 islands and the connecting open ocean throughout the Pacific Ocean (*37*) (Fig. 1A). The sampling strategy at each site was standardized and is detailed in (*38*). Briefly, 127 open ocean seawater samples (*ocean SW* in this study, *O-SRF* in dataset), 330 near island seawater samples (*island SW, I-SRFa*), 281 reef surface seawater samples (*reef SW*, *S-SRF*), 345 samples of seawater surrounding *Pocillopora* colonies (*coral SW*, *C-CSW*), and 219 reef sediment samples were collected (*sediment*), collectively referred to as environmental samples hereafter. At each site, a well-replicated set of coral samples were collected targeting the scleractinian branching morphotype *Pocillopora meandrina* (1,109 samples), massive *Porites lobata* (1,082 samples), and the hydrozoan firecoral *Millepora platyphylla* (652 samples). As hidden species diversity was previously identified (*43*), we therefore aggregated our results at the genus level (*Pocillopora*, *Porites*, and *Millepora*). In a complementary sampling approach, 80 additional coral colonies were randomly sampled at each island to capture overall coral diversity (CDIV) yielding 2,470 samples, of which 2,192 were available for analysis after quality filtering (*98*). These corals were taxonomically annotated to the lowest possible level based on a combination of *in situ* photographs and the 18S V9 marker (see (*38*) for detailed taxonomic assignment methods) resulting in 29 sampled lineages (*99*).

### DNA extraction, sequencing, and taxonomic assignment

Sample processing, DNA extraction, library preparation, and sequencing methods are detailed in (*40*). In summary, the sampling, processing, and extraction methods were adapted to each sample type (seawater, sediments, and corals), and we targeted the 18S V9 variable region with the primers 1389 F/1510 R (*100*). We processed raw reads with the nf-core/ampliseq workflow (*101*, *102*) version 2.8 (*103*) using default parameters unless stated otherwise. Primer sequences were removed (forward primer: 5’-*TTGTACACACCGCCC-3’*; reverse primer: 5’-*CCTTCYGCAGGTTCACCTAC-3’*), and all untrimmed sequences were discarded. Adapter and primer-free sequences were processed initially independently, but re-examined as one pool (pseudo-pooled) with DADA2 (*104*) to eliminate PhiX contamination, trim reads (forward reads at 90 bp and reverse reads at 90 bp, reads shorter than this were discarded), discard reads with > 2 expected errors, correct errors, merge read pairs, and remove polymerase chain reaction (PCR) chimeras. Following this, we filtered ASVs to remove those present in only a single sample or with fewer than three reads after dereplication. We assigned taxonomy using version v0.1.4 of an in-house Nextflow workflow using the IDTAXA algorithm from the R package DECIPHER version 2.22.0 (*105*) with R version 4.1.1 and the PR2 v.5.0.0 reference sequence database (*106*). This processing resulted in a total of 6,690 samples, 262,666 ASVS, and 3.42 x 10^9^ sequences. The full workflow is detailed in (*107*).

### Data processing

We applied several filtering steps to retain only target ASVs and samples. Target ASVs include protists and small metazoans, hereafter referred to as microeukaryotes. Specifically, we removed 11,646 ASVs assigned to Bacteria and Archaea, 113,944 ASVs with no close reference sequence (*no-hits*), 2 ASVs from PCR contamination, 3 mammalian ASVs, and 2,372 host-tissue ASVs (Anthozoan ASVs in scleractinian coral samples and coral surrounding seawater, as well as *Millepora* ASVs in *Millepora* samples). Supplementary Table 1 shows the number of ASVs and samples retained after each filtering step. We identified potential skin contaminants by analysing the most abundant ASVs in PCR controls. We removed only those ASVs that matched their reference sequence from the PR2 v.5.0.0 database (*106*) at 100%, which included the skin fungus *Malassezia globosa* and human DNA. We did not exclude Anthozoan ASVs from ocean, island, or reef seawater samples, as they accounted for only a small fraction of total sequences and may represent Anthozoan larvae (Supplementary Table 2). Most of the *no-hit* sequences occurred in sediments (25 % of total sequences), followed by seawater types (8-9 %), and were least abundant in corals (< 1.2 %; Supplementary Table 3). The combination of all filtering steps resulted in a total of 6,337 samples, 121,232 ASVS, and 1.26 x 10^9^ sequences. All data filtering and analyses were carried out in R (v4.5.1) with the packages *microbiome* (*108*) and *tidyverse* (*109*) for data handling and *ggplot2* (*110*)*, cowplot* (*111*), and *patchwork* (*112*) for plotting data.

### Alpha and beta diversity

For each sample type (ocean SW, island SW, reef SW, coral SW, sediments, *Millepora, Pocillopora* and *Porites*, and CDIV samples), we calculated the number of samples, total read counts, and minimum and maximum sequences per sample, as well as alpha diversity metrics of ASVs richness and Pielou’s evenness. We also identified the most abundant and the most prevalent ASVs. We repeated the same calculations for the pooled dataset across all sample types and compiled the results into an overview table.

We generated species accumulation curves to evaluate the increase in diversity with sample type (method = collector, 100 permutations). We also computed rarefaction curves to assess sampling effort, using a custom function that behaves as *rarefaction()* from the *vegan* package (*113*). The observed differences in sequencing depth across sample types were not attributed to technical variability in the sequencing process, but rather to the intrinsic biological properties of the sample, as coral samples consistently yielded substantially lower sequencing depth than sediment and water samples. Applying rarefaction would have resulted in the complete exclusion of most of *Porites* samples, thereby discarding a genuine biological signal rather than correcting for methodological bias. Samples were therefore normalised to relative abundance when indicated in the methods, following (*114*). Because assumptions of normality (Shapiro–Wilk test) and homogeneity of variances (Levene’s test) were not met (p < 0.05 in both cases, *car* package), we applied Wilcoxon rank-sum tests to compare observed ASV richness and Pielou’s evenness among sample types and adjusted p-values with the Benjamini–Hochberg procedure (FDR) (*rstatix*, *multcompView*, *ggplot2* packages).

We analysed beta diversity using non-metric multidimensional scaling (NMDS) in the *phyloseq* package (*115*). Because sampling protocols and DNA extraction methods differed among sample types, we avoided abundance-based comparisons of ASVs (*116*, *117*) and calculated community dissimilarities with the Jaccard index (presence–absence). We tested for differences in community composition with PERMANOVAs (*vegan* package) to evaluate group-level differences. We assessed the homogeneity of group dispersions using *betadisper* with Tukey’s HSD.

### Coral-associated eukaryome

To compare the abundances of microeukaryotic lineages among all sampled coral hosts (*Millepora, Pocillopora*, and *Porites*, plus CDIV samples), we removed octocoral samples (*Heliopora*) and coral taxa with fewer than five samples (including *Gardineroseris*, *Platygyra*, *Poritidae*, *Turbinaria*, *Lobophylliidae*, *Astrea*, *Coscinaraea*, *Herpolitha*, *Anthozona*, *Acanthastrea*, *Hydrozoa*, *Oxypora*, *Distichopora*, *Alveopora*, *Leptothecata*, *Euphyllia*, *Coeloseris*, *Physogyra*, *Stylocoeniella*, *Merulina*, *Acroporidae*, *Alcyonacea*, and *Coronatae*). We also excluded Symbiodiniaceae ASVs for this analysis, as their dominance would mask broader community patterns.

For each coral species, we summed ASV counts of the same taxonomic group, calculated the mean number of sequences per taxonomic group over all samples, and converted these values to relative abundance. We clustered corals by their eukaryotic community similarity using hierarchical clustering based on Bray–Curtis dissimilarities (*vegdist()* from the *vegan* package and *hclust()*). We visualised the dendrogram with the *ggdendro* package (*118*).

We extracted information on coral family and growth form from the Coral Trait Database (https://www.coraltraits.org/). We obtained data on life history traits and clades from (*62*), reproductive traits from (*63*), and information on *Millepora* from (*64*). Following this broad comparison over diverse coral lineages, only samples from the well-replicated genera *Millepora, Pocillopora,* and *Porites* were retained for further statistical inference as detailed below.

### Lineage counts

To summarise the diversity of microeukaryotic taxonomic lineages detected during the expedition, we grouped ASVs into their major monophyletic clades (hereafter referred to as lineages). For each lineage, we calculated total read counts, the number of ASVs, the total number of reference sequences used for taxonomic classification, and the proportion of ASVs assigned with different IDTAXA confidence ranges (50–60%, 60–70%, … 95–100%, and 100%) against PR2 (v5.0.0). We also analysed the correlation between the abundance and richness per lineage. We then assessed lineage distributions across samples in three ways. First, we calculated the relative distribution of sequences per lineage over the sample types. Second, we calculated the proportion of ASVs present in each sample type per lineage. Third, we quantified their biogeographical range as the number of Longhurst provinces (*41*) where each lineage occurred.

### Abundant taxa

In each sample type, we transformed ASV read counts into relative abundances within each sample. We then merged samples from the same site by calculating the mean relative abundance of each ASV across samples. Finally, we obtained lineage abundances per site by summing the relative abundances of all ASVs assigned to the same lineage, henceforth considered taxa. We retained only sites that contained both coral and either sediment or seawater samples (102 of 113 sites). In the coral and coral seawater samples, we removed Symbiodiniaceae ASVs prior to performing these calculations. We represented Symbiodiniaceae proportions separately, as the sum of all the ASV counts accounted for a large proportion of the sequences and would have hidden the underlying patterns. We repeated this procedure at the ASV level, without merging lineages, to determine whether the most abundant lineages were dominated by a large number of ASVs or by only a few.

### Shared, unique and prevalent ASVs

We visualised the overlap of ASVs among sample types using UpSet plots. We generated the intersection matrix using the *get_upset()* function from the *MicrobiotaProcess* package (*119*) and plotted it with the *UpSet()* function from *ComplexHeatmap* (*120*). For clarity, we displayed only intersections containing more than 200 ASVs. We calculated the prevalence of ASVs within each sample type, and extracted those ASVs that occurred in at least 50% of the samples from one of the three coral genera (*Millepora, Pocillopora*, and *Porites*) to identify highly prevalent ASVs.

### Network analysis for corals and seawater

We explored co-occurrence patterns of ASVs across reef-associated samples (ocean SW excluded) and within the three coral genera (*Millepora, Pocillopora,* and *Porites*). For this, we removed Symbiodiniaceae from the coral and coral SW samples, as they would link all host-associated ASVs and “force” the co-occurring patterns. We calculated the network using the FlashWeave algorithm (*121*), with the parameters heterogeneous = TRUE, sensitivity = TRUE, and a minimum observation threshold of 20 (default). We imported the results to R and retained only the positive associations to build the co-occurrence network and used the *igraph* package to explore the community structure (*122*). We detected modules of interconnected ASVs using the Louvain clustering algorithm *(cluster_louvain())*. We measured assortativity *(assortativity())*, to quantify whether ASVs preferably connect with others from the same taxonomic family, and assortativity degree *(assortativity_degree())* to assess if highly connected ASVs (hubs) tended to connect to other hubs. We calculated edge density *(edge_density())*, which represents the proportion of connections in the network from all possible ones, and the modularity *(vertex_connectivity())*, which evaluates the degree of network compartmentalisation. Visualisation was done with the *ggforce* package (*123*).

### Distance decay

We transformed all longitudes of the sampling sites to a 0-360° scale by adding 360° to negative values as sites spanned on opposite sides of the 180° meridian. Following this, we computed pairwise geographic distances between sites with the Haversine formula using the *distm()* function from *geosphere* package (*124*) and similarity matrices with Bray-Curtis dissimilarity. To test the relationship between community similarity and geographical distance, we performed Mantel tests for each sample type.

### Environmental drivers of microeukaryotic communities

We included composite SST variables, based on long-term remote sensing data (*125*) from 2002 to the sampling date, *in situ* measurements collected during the expedition, and modelled variables at each site (*126*). We also included latitude and longitude as explanatory variables (*127*). Full names, descriptions, and original datasets of the environmental variables are included in Supplementary Table 4. We retained only variables with coverage for more than 90% of samples. When variables were highly correlated (Pearson’s *r* > 0.7, VIF > 10), we selected a representative variable for downstream analyses (Supplementary Table 5).

We assessed the influence of environmental variables on community composition with a Bray–Curtis distance-based redundancy analysis (db-RDA) on scaled variables, using the *dbrda()* function in *vegan*. We selected db-RDA over CCA or RDA due to the high proportion of zeros in the dataset, typical in community matrices. We evaluated correlations between environmental variables and the community dissimilarity matrix with the *envfit()* function (*vegan)*, using 999 permutations. We further performed hierarchical clustering of environmental variables and coral hosts using Unweighted Pair Group Method with Arithmetic Mean (UPGMAM method *(hclust(.,method=average)*) on Euclidean distances computed from *envfit* r² values, to order the axes of the correlation heatmap.

### Comparison to OTU richness

Most of the global studies on marine eukaryotes that used amplicon sequencing, have relied on different bioinformatic pipelines inferring operational taxonomic units (OTUs) (*114*, *128*). In addition to our ASV-based analyses, we have calculated the total number of OTUs using the SWARM OTUs dataset (*129*) to facilitate a comparison of the diversity found in this expedition with other studies following the same filtering steps described in the *Data Filtering* section above.

## Acknowledgments

Special thanks to the Tara Ocean Foundation, the R/V Tara crew, and the *Tara* Pacific Expedition Participants (https://doi.org/10.5281/zenodo.3777760). We are keen to thank the commitment of the following institutions for their financial and scientific support that made this unique *Tara* Pacific Expedition possible: CNRS, PSL, CSM, EPHE, Genoscope, CEA, EMBL-EBI, Inserm, Université Côte d’Azur, ANR, agnès b., UNESCO-IOC, the Veolia Foundation, the Prince Albert II de Monaco Foundation, Région Bretagne, Billerudkorsnas, Amerisource Bergen Company, Lorient Agglomération, Oceans by Disney, L’Oréal, Biotherm, France Collectivités, Fonds Français pour l’Environnement Mondial (FFEM), Etienne Bourgois, and the Tara Ocean Foundation teams. *Tara* Pacific would not exist without the continuous support of the participating institutes. The authors also particularly thank Serge Planes, Denis Allemand, and the *Tara* Pacific consortium. We are grateful to the Roscoff Bioinformatics platform ABiMS (http://abims.sb-roscoff.fr), part of the Institut Français de Bioinformatique (ANR-11-INBS-0013) and BioGenouest network, for providing computing and storage resources. This work was supported by the de.NBI Cloud within the German Network for Bioinformatics Infrastructure (de.NBI) and ELIXIR-DE (Forschungszentrum Jülich and W-de.NBI-001, W-de.NBI-004, W-de.NBI-008, W-de.NBI-010, W-de.NBI-013, W-de.NBI-014, W-de.NBI-016, W-de.NBI-022). CRV and K-IM acknowledge funding from the Deutsche Forschungsgemeinschaft (DFG, German Research Foundation) project number 458901010. This is publication number #xx of the *Tara* Pacific Consortium.

## Data availability

All data used in this study is available in the TARA Pacific repository (https://zenodo.org/communities/tarapacific). Specifically, in this study we have used the following datasets: rDNA 18S V9 ASVs DADA2 v.1 (*107*) (https://zenodo.org/records/13837785), rDNA 18S V9 ASVs SWARM v.2 (*107*) (https://zenodo.org/records/10822169), CDIV cnidarian host taxonomic annotation v.1 (*98*) (https://zenodo.org/records/6327048), 18S-based coral host genetic analysis v.1 (*99*) (https://zenodo.org/records/4265266), samples provenance v.2 (*127*) (https://zenodo.org/records/6299409), environmental parameters at site level v.1 (*126*) (https://zenodo.org/records/6474974) and historical Sea Surface Temperature (SST) data and thermal stress indices v.20220317 (*125*)(https://zenodo.org/records/6499374).

Full code available at https://github.com/lauraDRH/TaraPacific_18S

## Author contributions

L.R.-H., M.Z., and the *Tara* Pacific Consortium Coordinators designed the research. M.Z. supervised the project. L.R.-H. and J.P. produced data. L.R.-H., N.H., C.L., C.R.V., B.C.C.H., and K.-I.M. curated data. L.R.-H. and N.H. analysed data. P.C.J., E.B., G.B., G.I., C.M., S.R., S.A., C.B., C.d.V., E.D., M.F., D.F., P.F., P.G., E.G., F.L., D.A.P.-G., S.P., S.R., S.S., O.T., M.M.R., R. T., R.V.T., C.R.V., P.W., D.Z., B.M.P., Q.C., M.Z., D.A. and S.P. managed the project, provided resources, or coordinated the field work. L.R.-H. and M.Z. drafted the manuscript with input from all authors. All of the authors reviewed the submitted manuscript and approved the final version.

## Competing interests

The authors declare no competing interests.

## Notes

### Competing Interest Statement

The authors have declared no competing interest.

## REFERENCES

1. M. L. Reaka-Kudla, “The global biodiversity of coral reefs: a comparison with rain forests” in Biodiversity II: Understanding and Protecting Our Biological Resources, M. L. Reaka-Kudla, D. E. Wilson, and E. O. Wilson, Ed. (Joseph Henry Press, 1996; https://books.google.com/books/about/Biodiversity_II.html?hl=&id=2-xRAgAAQBAJ), pp. 83–108.

2. F. Moberg, C. Folke, Ecological goods and services of coral reef ecosystems. Ecol. Econ. 29, 215–233 (1999).

3. C. Brown, E. Corcoran, P. Herkenrath, Marine and Coastal Ecosystems and Human Well-Being: A Synthesis Report Based on the Findings of the Millennium Ecosystem Assessment (UNEP, 2006; https://books.google.com/books/about/Marine_and_Coastal_Ecosystems_and_Human.html?hl=&id=7WoVAQAAIAAJ).

4. J. D. Reimer, R. S. Peixoto, S. W. Davies, N. Traylor-Knowles, M. L. Short, R. A. Cabral-Tena, J. A. Burt, I. Pessoa, A. T. Banaszak, R. S. Winters, T. Moore, V. Schoepf, D. Kaullysing, L. E. Calderon-Aguilera, G. Wörheide, S. Harding, V. Munbodhe, A. Mayfield, T. Ainsworth, T. Vardi, C. M. Eakin, M. S. Pratchett, C. R. Voolstra, The Fourth Global Coral Bleaching Event: Where do we go from here? Coral Reefs 43, 1121–1125 (2024).

5. P. Pearce-Kelly, C. Yesson, M. McField, A. I. Muñiz-Castillo, M. Soto, J. E. Arias-Gonzalez, K. Morgan, A. Bill-Weilandt, B. Kjerfve, C. E. Cornwall, L. Alvarez-Filip, M. Milkoreit, T. E. Lenton, R. M. Roman-Cuesta, “Warm-water coral reefs” in The Global Tipping Points Report 2025 (University of Exeter, UK, 2025), pp. 254–265.

6. B. E. Brown, Coral bleaching: causes and consequences. Coral Reefs 16, S129–S138 (1997).

7. D. G. Bourne, K. M. Morrow, N. S. Webster, Insights into the Coral Microbiome: Underpinning the Health and Resilience of Reef Ecosystems. Annual Review of Microbiology 70, 317–340 (2016).

8. J. R. Thompson, H. E. Rivera, C. J. Closek, M. Medina, Microbes in the coral holobiont: Partners through evolution, development, and ecological interactions. Frontiers in Cellular and Infection Microbiology 4 (2014).

9. P. E. Galand, H.-J. Ruscheweyh, G. Salazar, C. Hochart, N. Henry, B. C. C. Hume, P. H. Oliveira, A. Perdereau, K. Labadie, C. Belser, E. Boissin, S. Romac, J. Poulain, G. Bourdin, G. Iwankow, C. Moulin, E. J. Armstrong, D. A. Paz-García, M. Ziegler, S. Agostini, B. Banaigs, E. Boss, C. Bowler, C. de Vargas, E. Douville, M. Flores, D. Forcioli, P. Furla, E. Gilson, F. Lombard, S. Pesant, S. Reynaud, O. P. Thomas, R. Troublé, D. Zoccola, C. R. Voolstra, R. V. Thurber, S. Sunagawa, P. Wincker, D. Allemand, S. Planes, Diversity of the Pacific Ocean coral reef microbiome. Nature Communications 14, 3039 (2023).

10. I. Vanwonterghem, N. S. Webster, Coral Reef Microorganisms in a Changing Climate. iScience 23, 100972 (2020).

11. T. D. Ainsworth, A. J. Fordyce, E. F. Camp, The other microeukaryotes of the coral reef microbiome. Trends Microbiol. 25, 980–991 (2017).

12. J. Campo, D. Bass, P. J. Keeling, The eukaryome: Diversity and role of microeukaryotic organisms associated with animal hosts. Funct. Ecol. 34, 2045–2054 (2020).

13. J. Lukeš, C. R. Stensvold, K. Jirků-Pomajbíková, L. W. Parfrey, Are Human Intestinal Eukaryotes Beneficial or Commensals? PLOS Pathogens 11, e1005039 (2015).

14. C. Clerissi, S. Brunet, J. Vidal-Dupiol, M. Adjeroud, P. Lepage, L. Guillou, J.-M. Escoubas, E. Toulza, Protists within corals: The hidden diversity. Front. Microbiol. 9, 2043 (2018).

15. C. Ravindran, L. Irudayarajan, H. P. Raveendran, Possible beneficial interactions of ciliated protozoans with coral health and resilience. Appl Environ Microbiol 89, e0121723 (2023).

16. E. Kramarsky-Winter, M. Harel, N. Siboni, E. Ben Dov, I. Brickner, Y. Loya, A. Kushmaro, Identification of a protist-coral association and its possible ecological role. Mar. Ecol. Prog. Ser. 317, 67–73 (2006).

17. D. E. Morse, N. Hooker, A. N. C. Morse, R. A. Jensen, Control of larval metamorphosis and recruitment in sympatric agariciid corals. J. Exp. Mar. Bio. Ecol. 116, 193–217 (1988).

18. F. J. Pollock, R. McMinds, S. Smith, D. G. Bourne, B. L. Willis, M. Medina, R. V. Thurber, J. R. Zaneveld, Coral-associated bacteria demonstrate phylosymbiosis and cophylogeny. Nat. Commun. 9, 4921 (2018).

19. A. Hernandez-Agreda, W. Leggat, P. Bongaerts, C. Herrera, T. D. Ainsworth, Rethinking the coral microbiome: Simplicity exists within a diverse microbial biosphere. MBio 9 (2018).

20. R. S. Peixoto, C. R. Voolstra, Eds., Coral Reef Microbiome (Springer Nature Switzerland, Cham, 2025; 10.1007/978-3-031-76692-3).

21. F. Wiederkehr, L. Paoli, D. Richter, D. Racunica, H.-J. Ruscheweyh, M. Sperfeld, J. O’Brien, S. Miravet-Verde, A. B. Streiff, J. Ransome, C. Chepkirui, T. Priest, A. Sintsova, G. Salazar, K. S. I. Bistolas, T. Sawyer, K. Labadie, K.-I. Mayer, A. Perdereau, M. M. Reddy, C. Moulin, E. Boissin, G. Bourdin, J. Cailliau, G. Iwankow, J. Poulain, S. Romac, Tara Pacific Consortium Coordinators, C. de Vargas, J. M. Flores, P. Furla, E. Gilson, S. Pesant, S. Reynaud, D. Zoccola, S. Planes, D. Allemand, S. Agostini, C. Bowler, E. Douville, D. Forcioli, P. E. Galand, F. Lombard, P. H. Oliveira, O. P. Thomas, R. Vega Thurber, R. Troublé, C. R. Voolstra, P. Wincker, M. Ziegler, J. Piel, S. Sunagawa, Coral microbiomes as reservoirs of unknown genomic and biosynthetic diversity. Nature, doi: 10.1038/s41586-026-10159-6 (2026).

22. L. Muscatine, D. W. Goodall, The role of symbiotic algae in carbon and energy flux in reef corals. Coral Reefs 25, 75–87 (1992).

23. T. C. LaJeunesse, J. E. Parkinson, P. W. Gabrielson, H. J. Jeong, J. D. Reimer, C. R. Voolstra, S. R. Santos, Systematic Revision of Symbiodiniaceae Highlights the Antiquity and Diversity of Coral Endosymbionts. Curr. Biol. 28, 2570–2580.e6 (2018).

24. L. L. Blackall, B. Wilson, M. J. H. van Oppen, Coral-the world’s most diverse symbiotic ecosystem. Mol Ecol 24, 5330–5347 (2015).

25. M. J. Sweet, M. G. Séré, Ciliate communities consistently associated with coral diseases. J. Sea Res. 113, 119–131 (2016).

26. K. Priess, T. Le Campion-Alsumard, S. Golubic, F. Gadel, B. A. Thomassin, Fungi in corals: black bands and density-banding of Porites lutea and P. lobata skeleton. Mar. Biol. 136, 19–27 (2000).

27. W. K. Kwong, N. A. T. Irwin, V. Mathur, I. Na, N. Okamoto, M. J. A. Vermeij, P. J. Keeling, Taxonomy of the Apicomplexan Symbionts of Coral, including Corallicolida ord. nov., Reassignment of the Genus Gemmocystis, and Description of New Species Corallicola aquarius gen. nov. sp. nov. and Anthozoaphila gnarlus gen. nov. sp. nov. J Eukaryot Microbiol, e12852 (2021).

28. K. Tandon, M. M. Pasella, C. Iha, F. Ricci, J. Hu, C. J. O’Kelly, M. Medina, M. Kühl, H. Verbruggen, Every refuge has its price: Ostreobium as a model for understanding how algae can live in rock and stay in business. Semin Cell Dev Biol 134, 27–36 (2023).

29. D. Maggioni, R. Arrigoni, D. Seveso, P. Galli, M. L. Berumen, V. Denis, B. W. Hoeksema, D. Huang, F. Manca, D. Pica, S. Puce, J. D. Reimer, S. Montano, Evolution and biogeography of the Zanclea-Scleractinia symbiosis. Coral Reefs 41, 779–795 (2022).

30. C. Leboine, L. del Rio-Hortega, N. Henry, M. Zallio, A. M. Bonacolta, C. Belser, J.-M. Aury, C. R. Voolstra, B. C. C. Hume, A. Moussy, C. Moulin, E. Boissin, G. Bourdin, G. Iwankow, J. Poulain, S. Romac, Tara Pacific Consortium coordinators, J. del Campo, D. Allemand, S. Planes, M. Ziegler, P. Wincker, Q. Carradec, B. M. Porcel, Hidden Apicomplexan parasite diversity links coral and plankton microbiomes across reef seascapes, bioRxiv (2026)p. 2026.05.20.726672.

31. M. Zallio, C. Leboine, L. del Rio-Hortega, M. Ziegler, A. Moussy, C. Belser, F. Gavory, J.-M. Aury, D. Forcioli, P. Furla, T. Zamoum, K. Plichon, C. R. Voolstra, C. Moulin, E. Boissin, G. Bourdin, G. Iwankow, J. Poulain, S. Romac, Tara Pacific Consortium Coordinators, D. Allemand, S. Planes, P. Wincker, B. M. Porcel, Q. Carradec, Ocean-scale transcriptomic analysis of corallicolids (Apicomplexa) reveals their ubiquity and molecular interactions with *Pocillopora* corals, bioRxiv (2026). 10.64898/2026.05.20.726253.

32. J. del Campo, A. M. Bonacolta, B. A. Weiler, B. Knowles, A. Apprill, M. D. Fox, K. C. Wakeman, M. J. A. Vermeij, F. Rohwer, P. J. Keeling, Global diversity and distribution of coral-associated protists, bioRxiv (2026)p. 2026.01.22.700988.

33. B. Nogales, M. P. Lanfranconi, J. M. Piña-Villalonga, R. Bosch, Anthropogenic perturbations in marine microbial communities. FEMS Microbiol. Rev. 35, 275–298 (2011).

34. H. Dutta, A. Dutta, The microbial aspect of climate change. Energy Ecol. Environ. 1, 209–232 (2016).

35. D. A. Hutchins, F. Fu, Microorganisms and ocean global change. Nat. Microbiol. 2, 17058 (2017).

36. M. Ziegler, C. Arif, J. A. Burt, S. Dobretsov, C. Roder, T. C. LaJeunesse, C. R. Voolstra, Biogeography and molecular diversity of coral symbionts in the genus Symbiodinium around the Arabian Peninsula. J. Biogeogr. 44, 674–686 (2017).

37. S. Planes, D. Allemand, S. Agostini, B. Banaigs, E. Boissin, E. Boss, G. Bourdin, C. Bowler, E. Douville, J. M. Flores, D. Forcioli, P. Furla, P. E. Galand, J. F. Ghiglione, E. Gilson, F. Lombard, C. Moulin, S. Pesant, J. Poulain, S. Reynaud, S. Romac, M. B. Sullivan, S. Sunagawa, O. P. Thomas, R. Troublé, C. De Vargas, R. V. Thurber, C. R. Voolstra, P. Wincker, D. Zoccola, E. Armstrong, S. Audrain, J. M. Aury, B. Banaig, V. Barbe, C. Belser, E. Beraud, E. Bonnival, E. Bourgois, Q. Carradec, N. Cassar, N. R. Cohen, P. Conan, D. R. Cronin, O. Da Silva, N. Djerbi, J. R. Dolan, G. Dominguez Herta, J. Du, J. Filée, R. Friedrich, G. Gorsky, M. Guinther, N. Haëntjens, N. Henry, M. Hertau, C. Hochart, B. C. C. Hume, G. Iwankow, S. G. John, L. Karp-Boss, R. L. Kelly, Y. Kitano, G. Klinges, I. Koren, K. Labadie, J. Lancelot, N. Lang-Yona, J. Lê-Hoang, R. Lemee, Y. Lin, D. Marie, R. McMind, M. Miguel-Gordo, M. Trainic, D. Monmarche, Y. Mucherie, B. Noel, A. Ottaviani, L. Paoli, M. L. Pedrotti, C. Pogoreutz, M. Pujo-Pay, G. Reverdin, T. Röthig, E. Rottinger, A. Rouan, H. J. Ruscheweyh, G. Salazar, A. Vardi, R. Vega-Thunder, A. Zahed, T. Zamoum, The Tara Pacific expedition—A pan-ecosystemic approach of the “-omics” complexity of coral reef holobionts across the Pacific Ocean. [Preprint] (2019). 10.1371/journal.pbio.3000483.

38. F. Lombard, G. Bourdin, S. Pesant, S. Agostini, A. Baudena, E. Boissin, N. Cassar, M. Clampitt, P. Conan, O. Da Silva, C. Dimier, E. Douville, A. Elineau, J. Fin, J. M. Flores, J. F. Ghiglione, B. C. C. Hume, L. Jalabert, S. G. John, R. L. Kelly, I. Koren, Y. Lin, D. Marie, R. McMinds, Z. Mériguet, N. Metzl, D. A. Paz-García, M. L. Pedrotti, J. Poulain, M. Pujo-Pay, J. Ras, G. Reverdin, S. Romac, A. Rouan, E. Röttinger, A. Vardi, C. R. Voolstra, C. Moulin, G. Iwankow, B. Banaigs, C. Bowler, C. de Vargas, D. Forcioli, P. Furla, P. E. Galand, E. Gilson, S. Reynaud, S. Sunagawa, M. B. Sullivan, O. P. Thomas, R. Troublé, R. V. Thurber, P. Wincker, D. Zoccola, D. Allemand, S. Planes, E. Boss, G. Gorsky, Open science resources from the Tara Pacific expedition across coral reef and surface ocean ecosystems. Sci. Data 10, 1–25 (2023).

39. S. Planes, D. Allemand, Insights and achievements from the Tara Pacific expedition. Nature Research [Preprint] (2023). 10.1038/s41467-023-38896-6.

40. C. Belser, J. Poulain, K. Labadie, F. Gavory, A. Alberti, J. Guy, Q. Carradec, C. Cruaud, C. Da Silva, S. Engelen, P. Mielle, A. Perdereau, G. Samson, S. Gas, Genoscope Technical Team, C. R. Voolstra, P. E. Galand, J. M. Flores, B. C. C. Hume, G. Perna, M. Ziegler, H.-J. Ruscheweyh, E. Boissin, S. Romac, G. Bourdin, G. Iwankow, C. Moulin, D. A. Paz García, S. Agostini, B. Banaigs, E. Boss, C. Bowler, C. de Vargas, E. Douville, D. Forcioli, P. Furla, E. Gilson, F. Lombard, S. Pesant, S. Reynaud, S. Sunagawa, O. P. Thomas, R. Troublé, R. V. Thurber, D. Zoccola, C. Scarpelli, E. K. Jacoby, P. H. Oliveira, J.-M. Aury, D. Allemand, S. Planes, P. Wincker, Integrative omics framework for characterization of coral reef ecosystems from the Tara Pacific expedition. Sci. Data 10, 326 (2023).

41. A. R. Longhurst, Ecological Geography of the Sea (Academic Press, San Diego, CA, ed. 2, 2006; 10.1016/b978-0-12-455521-1.x5000-1).

42. J. Arañó-Ansola, I. Galán-Luque, M. Domènech, L. Rico-Martín, E. R. R. Moody, M. Suresh, D. Vaulot, J. del Campo, M. Giacomelli, J. Lozano-Fernandez, Evaluating 18S phylogenetic placement accuracy to uncover hidden diversity in early branching animals, bioRxiv (2025)p. 2025.12.09.693319.

43. C. R. Voolstra, B. C. C. Hume, E. J. Armstrong, G. Mitushasi, B. Porro, N. Oury, S. Agostini, E. Boissin, J. Poulain, Q. Carradec, D. A. Paz-García, D. Zoccola, H. Magalon, C. Moulin, G. Bourdin, G. Iwankow, S. Romac, B. Banaigs, E. Boss, C. Bowler, C. de Vargas, E. Douville, M. Flores, P. Furla, P. E. Galand, E. Gilson, F. Lombard, S. Pesant, S. Reynaud, M. B. Sullivan, S. Sunagawa, O. P. Thomas, R. Troublé, R. V. Thurber, P. Wincker, S. Planes, D. Allemand, D. Forcioli, Disparate genetic divergence patterns in three corals across a pan-Pacific environmental gradient highlight species-specific adaptation. NPJ Biodivers 2, 15 (2023).

44. N. Henry, J. Poulain, C. de Vargas, Tara Oceans (2009-2013) rDNA 18S V9 ASV table (DADA2) with nf-core/ampliseq 2.13, Zenodo (2026); 10.5281/ZENODO.18153978.

45. F. Aslani, S. Geisen, D. Ning, L. Tedersoo, M. Bahram, Towards revealing the global diversity and community assembly of soil eukaryotes. Ecol. Lett. 25, 65–76 (2022).

46. S. Y. Moon-van der Staay, R. De Wachter, D. Vaulot, Oceanic 18S rDNA sequences from picoplankton reveal unsuspected eukaryotic diversity. Nature 409, 607–610 (2001).

47. L. R. Thompson, J. G. Sanders, D. McDonald, A. Amir, J. Ladau, K. J. Locey, R. J. Prill, A. Tripathi, S. M. Gibbons, G. Ackermann, J. A. Navas-Molina, S. Janssen, E. Kopylova, Y. Vázquez-Baeza, A. González, J. T. Morton, S. Mirarab, Z. Zech Xu, L. Jiang, M. F. Haroon, J. Kanbar, Q. Zhu, S. Jin Song, T. Kosciolek, N. A. Bokulich, J. Lefler, C. J. Brislawn, G. Humphrey, S. M. Owens, J. Hampton-Marcell, D. Berg-Lyons, V. McKenzie, N. Fierer, J. A. Fuhrman, A. Clauset, R. L. Stevens, A. Shade, K. S. Pollard, K. D. Goodwin, J. K. Jansson, J. A. Gilbert, R. Knight, Earth Microbiome Project Consortium, A communal catalogue reveals Earth’s multiscale microbial diversity. Nature 551, 457–463 (2017).

48. S. M. Bower, R. B. Carnegie, B. Goh, S. R. Jones, G. J. Lowe, M. W. Mak, Preferential PCR amplification of parasitic protistan small subunit rDNA from metazoan tissues. J Eukaryot Microbiol 51, 325–332 (2004).

49. A. Góes-Neto, V. R. Marcelino, H. Verbruggen, F. F. da Silva, F. Badotti, Biodiversity of endolithic fungi in coral skeletons and other reef substrates revealed with 18S rDNA metabarcoding. Coral Reefs 39, 229–238 (2020).

50. A. M. Bonacolta, B. A. Weiler, T. Porta-Fitó, M. Sweet, P. Keeling, J. del Campo, Beyond the Symbiodiniaceae: diversity and role of microeukaryotic coral symbionts. Coral Reefs 42, 567–577 (2023).

51. K. M. Morrow, M. S. Pankey, M. P. Lesser, Community structure of coral microbiomes is dependent on host morphology. Microbiome 10, 113 (2022).

52. J. Liang, K. Yu, Y. Wang, X. Huang, W. Huang, Z. Qin, Z. Pan, Q. Yao, W. Wang, Z. Wu, Distinct bacterial communities associated with massive and branching scleractinian corals and potential linkages to coral susceptibility to thermal or cold stress. Front. Microbiol. 8, 979 (2017).

53. S. Schmitt, P. Tsai, J. Bell, J. Fromont, M. Ilan, N. Lindquist, T. Perez, A. Rodrigo, P. J. Schupp, J. Vacelet, N. Webster, U. Hentschel, M. W. Taylor, Assessing the complex sponge microbiota: core, variable and species-specific bacterial communities in marine sponges. ISME J 6, 564–576 (2012).

54. J. B. Clayton, A. Gomez, K. Amato, D. Knights, D. A. Travis, R. Blekhman, R. Knight, S. Leigh, R. Stumpf, T. Wolf, K. E. Glander, F. Cabana, T. J. Johnson, The gut microbiome of nonhuman primates: Lessons in ecology and evolution. Am J Primatol 80, e22867 (2018).

55. H. E. Epstein, T. Brown, A. O. Akinrinade, R. McMinds, F. J. Pollock, D. Sonett, S. Smith, D. G. Bourne, C. S. Carpenter, R. Knight, B. L. Willis, M. Medina, J. B. Lamb, R. V. Thurber, J. R. Zaneveld, Evidence for microbially-mediated tradeoffs between growth and defense throughout coral evolution. *Anim*. Microbiome 7, 1 (2025).

56. B. W. Hoeksema, S. E. T. Van der Meij, C. H. J. M. Fransen, The mushroom coral as a habitat. J. Mar. Biol. Assoc. U. K. 92, 647–663 (2012).

57. Y. R. Cheng, A. B. Mayfield, P. J. Meng, C. F. Dai, R. Huys, Copepods associated with scleractinian corals: a worldwide checklist and a case study of their impact on the reef-building coral Pocillopora damicornis (Linnaeus, 1758) (Pocilloporidae). Zootaxa 4174, 291–345 (2016).

58. K. Y. Inagaki, G. O. Longo, Revisiting 20 years of coral–algal interactions: global patterns and knowledge gaps. Coral Reefs 43, 899–917 (2024).

59. W. Gong, A. Marchetti, Estimation of 18S gene copy number in marine eukaryotic plankton using a next-generation sequencing approach. Front. Mar. Sci. 6, 453814 (2019).

60. P. D. Lamb, E. Hunter, J. K. Pinnegar, S. Creer, R. G. Davies, M. I. Taylor, How quantitative is metabarcoding: A meta-analytical approach. Mol. Ecol. 28, 420–430 (2019).

61. E. Hehenberger, D. V. Tikhonenkov, M. Kolisko, J. Del Campo, A. S. Esaulov, A. P. Mylnikov, P. J. Keeling, Novel predators reshape holozoan phylogeny and reveal the presence of a two-component signaling system in the ancestor of animals. Curr. Biol. 27, 2043–2050.e6 (2017).

62. E. S. Darling, L. Alvarez-Filip, T. A. Oliver, T. R. McClanahan, I. M. Côté, D. Bellwood, Evaluating life-history strategies of reef corals from species traits. Ecol Lett 15, 1378–1386 (2012).

63. A. H. Baird, J. R. Guest, B. L. Willis, Systematic and biogeographical patterns in the reproductive biology of scleractinian corals. Annual Review of Ecology, Evolution, and Systematics 40, 551–571 (2009).

64. J. B. Lewis, Biology and ecology of the hydrocoral millepora on coral reefs. Adv. Mar. Biol. 50, 1–55 (2006).

65. C. Iha, K. E. Dougan, J. A. Varela, V. Avila, C. J. Jackson, K. A. Bogaert, Y. Chen, L. M. Judd, R. Wick, K. E. Holt, M. M. Pasella, F. Ricci, S. I. Repetti, M. Medina, V. R. Marcelino, C. X. Chan, H. Verbruggen, Genomic adaptations to an endolithic lifestyle in the coral-associated alga Ostreobium. Curr Biol 31, 1393–1402.e5 (2021).

66. W. K. Kwong, J. Del Campo, V. Mathur, M. J. A. Vermeij, P. J. Keeling, A widespread coral-infecting apicomplexan with chlorophyll biosynthesis genes. Nature 568, 103–107 (2019).

67. A. Peterson, S. Patton, E. R. Schmeltzer, C. G. B. Grupstra, L. I. Howe-Kerr, J. G. Klinges, R. L. Maher, A. Messyasz, S. Seabrook, A. R. Thurber, A. M. S. Correa, R. L. Vega Thurber, Apicomplexan and non-metazoan microeukaryotes in the thermosensitive reef-building coral Acropora hyacinthus shift in abundance throughout an extreme coral bleaching event. Front. Mar. Sci. 12, 1626071 (2025).

68. P. J. B. Scott, M. J. Risk, The effect of Lithophaga (Bivalvia: Mytilidae) boreholes on the strength of the coral Porites lobata. Coral Reefs 7, 145–151 (1988).

69. P.-L. Stenger, A. Tribollet, F. Guilhaumon, P. Cuet, G. Pennober, P. Jourand, A Multimarker Approach to Identify Microbial Bioindicators for Coral Reef Health Monitoring-Case Study in La Réunion Island. Microb Ecol 87, 179 (2025).

70. P. Hallock, B. H. Lidz, E. M. Cockey-Burkhard, K. B. Donnelly, Foraminifera as bioindicators in coral reef assessment and monitoring: The foram index. Environ. Monit. Assess. 81, 221–238 (2003).

71. R. Logares, I. M. Deutschmann, P. C. Junger, C. R. Giner, A. K. Krabberød, T. S. B. Schmidt, L. Rubinat-Ripoll, M. Mestre, G. Salazar, C. Ruiz-González, M. Sebastián, C. de Vargas, S. G. Acinas, C. M. Duarte, J. M. Gasol, R. Massana, Disentangling the mechanisms shaping the surface ocean microbiota. Microbiome 8, 55 (2020).

72. L. Weber, P. Gonzalez-Díaz, M. Armenteros, A. Apprill, The coral ecosphere: A unique coral reef habitat that fosters coral–microbial interactions. Limnology and Oceanography 64, 2373–2388 (2019).

73. K. Walsh, J. M. Haggerty, M. P. Doane, J. J. Hansen, M. M. Morris, A. P. B. Moreira, L. de Oliveira, L. Leomil, G. D. Garcia, F. Thompson, E. A. Dinsdale, Aura-biomes are present in the water layer above coral reef benthic macro-organisms. PeerJ 2017, e3666 (2017).

74. W. C. Million, C. R. Voolstra, G. Perna, G. Puntin, K. Rowe, M. Ziegler, Resolving Symbiodiniaceae Diversity Across Coral Microhabitats and Reef Niches. Environ Microbiol 27, e70065 (2025).

75. F. Wiederkehr, K. E. Engelhardt, J. Vetter, H.-J. Ruscheweyh, G. Salazar, J. O’Brien, T. Priest, M. Ziegler, S. Sunagawa, Host-level biodiversity shapes the dynamics and networks within the coral reef microbiome. ISME Commun 5, ycaf097 (2025).

76. J. C. W. Liu, J. T. Høeg, B. K. K. Chan, How do coral barnacles start their life in their hosts? Biol Lett 12 (2016).

77. J. R. Griffiths, M. Kadin, F. J. A. Nascimento, T. Tamelander, A. Törnroos, S. Bonaglia, E. Bonsdorff, V. Brüchert, A. Gårdmark, M. Järnström, J. Kotta, M. Lindegren, M. C. Nordström, A. Norkko, J. Olsson, B. Weigel, R. Žydelis, T. Blenckner, S. Niiranen, M. Winder, The importance of benthic-pelagic coupling for marine ecosystem functioning in a changing world. Glob Chang Biol 23, 2179–2196 (2017).

78. B. J. Baker, K. E. Appler, X. Gong, New Microbial Biodiversity in Marine Sediments. Ann Rev Mar Sci 13, 161–175 (2021).

79. E. Capo, D. Debroas, F. Arnaud, I. Domaizon, Is Planktonic Diversity Well Recorded in Sedimentary DNA? Toward the Reconstruction of Past Protistan Diversity. Microb Ecol 70, 865–875 (2015).

80. D. P. Tittensor, C. Mora, W. Jetz, H. K. Lotze, D. Ricard, E. V. Berghe, B. Worm, Global patterns and predictors of marine biodiversity across taxa. Nature 466, 1098–1101 (2010).

81. A. H. Altieri, K. B. Gedan, Climate change and dead zones. Glob Chang Biol 21, 1395–1406 (2015).

82. M. Voss, H. W. Bange, J. W. Dippner, J. J. Middelburg, J. P. Montoya, B. Ward, The marine nitrogen cycle: recent discoveries, uncertainties and the potential relevance of climate change. Philos Trans R Soc Lond B Biol Sci 368, 20130121 (2013).

83. Y. Wang, J. B. Kajtar, L. V. Alexander, G. S. Pilo, N. J. Holbrook, Understanding the changing nature of marine cold-spells. Geophys. Res. Lett. 49 (2022).

84. Y. Yao, C. Wang, Marine heatwaves and cold-spells in global coral reef zones. Prog. Oceanogr. 209, 102920 (2022).

85. T. K. Watanabe, S. Ito, A. U. Nurhidayati, S. Y. Cahyarini, M. Pfeiffer, Coral reefs in the Indonesian Seas threatened by heat and cold stress. Geophys. Res. Lett. 53, e2025GL121003 (2026).

86. S. Chaffron, E. Delage, M. Budinich, D. Vintache, N. Henry, C. Nef, M. Ardyna, A. A. Zayed, P. C. Junger, P. E. Galand, C. Lovejoy, A. E. Murray, H. Sarmento, Tara Oceans coordinators, S. G. Acinas, M. Babin, D. Iudicone, O. Jaillon, E. Karsenti, P. Wincker, L. Karp-Boss, M. B. Sullivan, C. Bowler, C. de Vargas, D. Eveillard, Environmental vulnerability of the global ocean epipelagic plankton community interactome. Sci. Adv. 7, eabg1921 (2021).

87. S. L. Coles, P. L. Jokiel, Synergistic effects of temperature, salinity and light on the hermatypic coral Montipora verrucosa. Mar. Biol. 49, 187–195 (1978).

88. D. Seveso, S. Montano, G. Strona, I. Orlandi, P. Galli, M. Vai, Exploring the effect of salinity changes on the levels of Hsp60 in the tropical coral Seriatopora caliendrum. Mar Environ Res 90, 96–103 (2013).

89. C. Ferrier-Pagès, J. P. Gattuso, J. Jaubert, Effect of small variations in salinity on the rates of photosynthesis and respiration of the zooxanthellate coral Stylophora pistillata. Mar. Ecol. Prog. Ser. 181, 309–314 (1999).

90. N. Rädecker, S. Escrig, J. E. Spangenberg, C. R. Voolstra, A. Meibom, Coupled carbon and nitrogen cycling regulates the cnidarian-algal symbiosis. Nat Commun 14, 6948 (2023).

91. T. Xiang, E. Lehnert, R. E. Jinkerson, S. Clowez, R. G. Kim, J. C. DeNofrio, J. R. Pringle, A. R. Grossman, Symbiont population control by host-symbiont metabolic interaction in Symbiodiniaceae-cnidarian associations. Nat Commun 11, 108 (2020).

92. C. Pogoreutz, N. Rädecker, A. Cárdenas, A. Gärdes, C. Wild, C. R. Voolstra, Dominance of *Endozoicomonas* bacteria throughout coral bleaching and mortality suggests structural inflexibility of the *Pocillopora verrucosa* microbiome. Ecol. Evol. 8, 2240–2252 (2018).

93. M. J. Neave, R. Rachmawati, L. Xun, C. T. Michell, D. G. Bourne, A. Apprill, C. R. Voolstra, Differential specificity between closely related corals and abundant *Endozoicomonas* endosymbionts across global scales. ISME J. 11, 186–200 (2016).

94. S. R. Proulx, D. E. L. Promislow, P. C. Phillips, Network thinking in ecology and evolution. Elsevier Ltd [Preprint] (2005). 10.1016/j.tree.2005.04.004.

95. O. Artime, M. Grassia, M. De Domenico, J. P. Gleeson, H. A. Makse, G. Mangioni, M. Perc, F. Radicchi, Robustness and resilience of complex networks. Nat. Rev. Phys. 6, 114–131 (2024).

96. D. Berry, S. Widder, Deciphering microbial interactions and detecting keystone species with co-occurrence networks. Front. Microbiol. 5, 219 (2014).

97. A. M. Martín González, B. Dalsgaard, J. M. Olesen, Centrality measures and the importance of generalist species in pollination networks. Ecol. Complex. 7, 36–43 (2010).

98. B. C. C. Hume, Y. Kitano, H. Fukami, S. Agostini, J. Poulain, S. Pesant, C. Belser, H.-J. Ruscheweyh, D. A. Paz Garcia, E. Armstrong, Q. Clayssen, N. Henry, G. Klinges, R. McMinds, L. Paoli, C. Pogoreutz, G. Salazar, M. Ziegler, C. Moulin, E. Boissin, G. Bourdin, G. Iwankow, S. Romac, B. Banaigs, E. Boss, C. Bowler, C. de Vargas, E. Douville, J. M. Flores, P. Furla, P. E. Galand, E. Gilson, F. Lombard, S. Reynaud, O. Thomas, R. Troublé, R. Vega Thurber, D. Zoccola, S. Planes, D. Allemand, S. Sunagawa, P. Wincker, D. Forcioli, C. R. Voolstra, TARA Pacific CDIV cnidarian host taxonomic annotation release version 1_1, Zenodo (2022); 10.5281/ZENODO.6327047.

99. B. C. C. Hume, J. Poulain, S. Pesant, C. Belser, H.-J. Ruscheweyh, E. Boissin, E. Armstrong, Q. Clayssen, N. Henry, G. Klinges, R. McMinds, L. Paoli, C. Pogoreutz, G. Salazar, M. Ziegler, C. Moulin, G. Bourdin, G. Iwankow, S. Romac, S. Agostini, B. Banaigs, E. Boss, C. Bowler, C. de Vargas, E. Douville, M. Flores, P. Furla, P. E. Galand, E. Gilson, F. Lombard, S. Reynaud, M. B. Sullivan, S. Sunagawa, O. Thomas, R. Troublé, R. Vega Thurber, D. Zoccola, S. Planes, D. Allemand, D. Forcioli, P. Wincker, C. R. Voolstra, Tara Pacific 18S-based coral host genetic analysis data release version 1, Zenodo (2020); 10.5281/ZENODO.4265266.

100. L. A. Amaral-Zettler, E. A. McCliment, H. W. Ducklow, S. M. Huse, A method for studying protistan diversity using massively parallel sequencing of V9 hypervariable regions of small-subunit ribosomal RNA genes. PLoS One 4, e6372 (2009).

101. D. Straub, N. Blackwell, A. Langarica-Fuentes, A. Peltzer, S. Nahnsen, S. Kleindienst, Interpretations of Environmental Microbial Community Studies Are Biased by the Selected 16S rRNA (Gene) Amplicon Sequencing Pipeline. Front Microbiol 11, 550420 (2020).

102. P. A. Ewels, A. Peltzer, S. Fillinger, H. Patel, J. Alneberg, A. Wilm, M. U. Garcia, P. Di Tommaso, S. Nahnsen, The nf-core framework for community-curated bioinformatics pipelines. Nat Biotechnol 38, 276–278 (2020).

103. D. Straub, J. Tångrot, A. Peltzer, D. Lundin, emnilsson, N.-C. Bot, Sateesh_Peri, A. Bennett, J. Sundh, DiegoBrambilla, A. Peer, T. E., L. Manoharan, M. U. Garcia, S. Minot, PhilPalmer, H. Patel, V. Malladi, Matthew, G. Gabernet, D. Clayton, P. Ewels, C. Davenport, dariader, K. Menden, Nf-Core/ampliseq: Ampliseq *Version 2.8.0* (Zenodo, 2024; https://zenodo.org/doi/10.5281/zenodo.10519258).

104. B. J. Callahan, P. J. McMurdie, M. J. Rosen, A. W. Han, A. J. A. Johnson, S. P. Holmes, DADA2: High-resolution sample inference from Illumina amplicon data. Nat. Methods 13, 581–583 (2016).

105. E. Wright, Using DECIPHER v2.0 to analyze big biological sequence data in R. R J. 8, 352 (2016).

106. L. Guillou, D. Bachar, S. Audic, D. Bass, C. Berney, L. Bittner, C. Boutte, G. Burgaud, C. de Vargas, J. Decelle, J. Del Campo, J. R. Dolan, M. Dunthorn, B. Edvardsen, M. Holzmann, W. H. C. F. Kooistra, E. Lara, N. Le Bescot, R. Logares, F. Mahé, R. Massana, M. Montresor, R. Morard, F. Not, J. Pawlowski, I. Probert, A.-L. Sauvadet, R. Siano, T. Stoeck, D. Vaulot, P. Zimmermann, R. Christen, The Protist Ribosomal Reference database (PR2): a catalog of unicellular eukaryote small sub-unit rRNA sequences with curated taxonomy. Nucleic Acids Res 41, D597–604 (2013).

107. N. Henry, F. Mahé, C. Berney, J. Poulain, S. Romac, G. Bourdin, H.-J. Ruscheweyh, G. Salazar, C. Belser, Q. Clayssen, B. Hume, E. Boissin, P. E. Galand, S. Pesant, F. Lombard, E. Armstrong, N. Lang-Yona, R. McMinds, R. Vega Thurber, C. Moulin, S. Agostini, B. Banaigs, E. Boss, C. Bowler, E. Douville, J. M. Flores, D. Forcioli, P. Furla, E. Gilson, S. Reynaud, M. Sullivan, O. P. Thomas, R. Troublé, D. Zoccola, S. Planes, D. Allemand, S. Sunagawa, C. R. Voolstra, P. Wincker, C. de Vargas, rDNA 18S V9 ASVs (DADA2) from the Tara Pacific Expedition, Zenodo (2024); 10.5281/ZENODO.13837784.

108. L. Lahti, S. Shetty, Microbiome r package: Tools for microbiome analysis in r. Bioconductor, doi: https://github.com/microbiome/microbiome (2017).

109. H. Wickham, M. Averick, J. Bryan, W. Chang, L. McGowan, R. François, G. Grolemund, A. Hayes, L. Henry, J. Hester, M. Kuhn, T. Pedersen, E. Miller, S. Bache, K. Müller, J. Ooms, D. Robinson, D. Seidel, V. Spinu, K. Takahashi, D. Vaughan, C. Wilke, K. Woo, H. Yutani, Welcome to the tidyverse. J. Open Source Softw. 4, 1686 (2019).

110. H. Wickham, W. Chang, L. Henry, T. L. Pedersen, K. Takahashi, C. Wilke, K. Woo, H. Yutani, D. Dunnington, T. van den Brand, Ggplot2: Create elegant data visualisations using the grammar of graphics, The R Foundation (2007); 10.32614/cran.package.ggplot2.

111. C. O. Wilke, cowplot: Streamlined Plot Theme and Plot Annotations for “ggplot2,” The R Foundation (2015); 10.32614/cran.package.cowplot.

112. T. L. Pedersen, patchwork: The Composer of Plots, The R Foundation (2019); 10.32614/cran.package.patchwork.

113. J. Oksanen, G. Simpson, F. Blanchet, R. Kindt, P. Legendre, P. Minchin, R. O’Hara, P. Solymos, M. Stevens, E. Szoecs, H. Wagner, M. Barbour, M. Bedward, B. Bolker, D. Borcard, G. Carvalho, M. Chirico, M. De Caceres, S. Durand, H. Evangelista, R. FitzJohn, M. Friendly, B. Furneaux, G. Hannigan, M. Hill, L. Lahti, D. McGlinn, M. Ouellette, E. Ribeiro Cunha, T. Smith, A. Stier, C. Ter Braak, J. Weedon, vegan: Community Ecology Package. R package version 2.6–5. doi: https://github.com/vegandevs/vegan. (2023).

114. C. De Vargas, S. Audic, N. Henry, J. Decelle, F. Mahé, R. Logares, E. Lara, Ć. Berney, N. Le Bescot, I. Probert, M. Carmichael, J. Poulain, S. Romac, S. Colin, J. M. Aury, L. Bittner, S. Chaffron, M. Dunthorn, S. Engelen, O. Flegontova, L. Guidi, A. Horák, O. Jaillon, G. Lima-Mendez, J. Lukeš, S. Malviya, R. Morard, M. Mulot, E. Scalco, R. Siano, F. Vincent, A. Zingone, C. Dimier, M. Picheral, S. Searson, S. Kandels-Lewis, S. G. Acinas, P. Bork, C. Bowler, G. Gorsky, N. Grimsley, P. Hingamp, D. Iudicone, F. Not, H. Ogata, S. Pesant, J. Raes, M. E. Sieracki, S. Speich, L. Stemmann, S. Sunagawa, J. Weissenbach, P. Wincker, E. Karsenti, E. Boss, M. Follows, L. Karp-Boss, U. Krzic, E. G. Reynaud, C. Sardet, M. B. Sullivan, D. Velayoudon, Ocean plankton. Eukaryotic plankton diversity in the sunlit ocean. Science 348 (2015).

115. P. J. McMurdie, S. Holmes, Phyloseq: An R Package for Reproducible Interactive Analysis and Graphics of Microbiome Census Data. PLoS ONE 8 (2013).

116. A. Dopheide, D. Xie, T. R. Buckley, A. J. Drummond, R. D. Newcomb, Impacts of DNA extraction and PCR on DNA metabarcoding estimates of soil biodiversity. Methods Ecol. Evol. 10, 120–133 (2019).

117. V. Vasselon, I. Domaizon, F. Rimet, M. Kahlert, A. Bouchez, Application of high-throughput sequencing (HTS) metabarcoding to diatom biomonitoring: Do DNA extraction methods matter? Freshw. Sci. 36, 162–177 (2017).

118. A. de Vries, B. D. Ripley, ggdendro: Create Dendrograms and Tree Diagrams Using “ggplot2,” The R Foundation (2013); 10.32614/cran.package.ggdendro.

119. S. Xu, L. Zhan, W. Tang, Q. Wang, Z. Dai, L. Zhou, T. Feng, M. Chen, T. Wu, E. Hu, G. Yu, MicrobiotaProcess: A comprehensive R package for deep mining microbiome. Innovation (Camb.) 4, 100388 (2023).

120. Z. Gu, ComplexHeatmap (Bioconductor, 2017; https://bioconductor.org/packages/ComplexHeatmap).

121. J. Tackmann, J. F. Matias Rodrigues, C. von Mering, Rapid inference of direct interactions in large-scale ecological networks from heterogeneous microbial sequencing data. Cell Syst. 9, 286–296.e8 (2019).

122. G. Csárdi, T. Nepusz, V. Traag, S. Horvát, F. Zanini, D. Noom, K. Müller, D. Schoch, M. Salmon, Igraph: Network analysis and visualization, The R Foundation (2006); 10.32614/cran.package.igraph.

123. T. L. Pedersen, ggforce: Accelerating “ggplot2,” The R Foundation (2016); 10.32614/cran.package.ggforce.

124. R. J. Hijmans, geosphere: Spherical Trigonometry, The R Foundation (2010); 10.32614/cran.package.geosphere.

125. G. Bourdin, F. Lombard, E. Boss, G. Gorsky, S. Pesant, C. R. Voolstra, C. Moulin, E. Boissin, G. Iwankow, J. Poulain, S. Romac, S. Agostini, B. Banaigs, C. Bowler, C. de Vargas, E. Douville, J. M. Flores, D. Forcioli, P. Furla, P. E. Galand, E. Gilson, S. Reynaud, M. B. Sullivan, S. Sunagawa, O. Thomas, R. Troublé, R. Vega Thurber, P. Wincker, D. Zoccola, S. Planes, D. Allemand, Tara Pacific Consortium, Historical Sea Surface Temperature (SST) data and thermal stress indices of the Tara Pacific Expedition’s coral reef sampling sites, from May 1st 2002 to August 31st 2018, Zenodo (2022); 10.5281/ZENODO.6499373.

126. Bourdin, Lombard, Boss, Douville, Flores, Cassar, Cohen, Dimier, Fin, Gorsky, John, Kelly, Koren, Lin, Marie, Metzl, Pujo-Pay, Ras, Reverdin, Vardi, Conan, Ghiglione, Moulin, Boissin, Iwankow, Poulain, Romac, Agostini, Banaigs, Bowler, D. Vargas, Forcioli, Furla, P. E. Galand, E. Gilson, S. Pesant, S. Reynaud, M. B. Sullivan, S. Sunagawa, O. Thomas, R. Troublé, R. Vega Thurber, C. R. Voolstra, P. Wincker, D. Zoccola, D. Allemand, S. Planes, Environmental context observed during the Tara Pacific Expedition 2016-2018, simplified version at site level, Zenodo (2022); 10.5281/ZENODO.6474973.

127. S. Pesant, F. Lombard, G. Bourdin, J. Poulain, E. Petit, E. Boss, N. Cassar, N. R. Cohen, C. Dimier, E. Douville, J. M. Flores, G. Gorsky, B. C. C. Hume, S. G. John, R. L. Kelly, Y. Lin, D. Marie, M.-L. Pedrotti, M. Pujo Pay, J. Ras, G. Reverdin, H.-J. Ruscheweyh, A. Vardi, C. R. Voolstra, C. Moulin, E. Boissin, G. Iwankow, S. Romac, S. Agostini, B. Banaigs, C. Bowler, C. De Vargas, D. Forcioli, P. Furla, P. E. Galand, E. Gilson, S. Reynaud, S. M. B., S. Sunagawa, O. Thomas, R. Troublé, R. Vega Thurber, P. Wincker, D. Zoccola, S. Planes, D. Allemand, Tara Pacific samples provenance and environmental context - version 2, Zenodo (2020); 10.5281/ZENODO.4068292.

128. F. M. Ibarbalz, N. Henry, M. C. Brandão, S. Martini, G. Busseni, H. Byrne, L. P. Coelho, H. Endo, J. M. Gasol, A. C. Gregory, F. Mahé, J. Rigonato, M. Royo-Llonch, G. Salazar, I. Sanz-Sáez, E. Scalco, D. Soviadan, A. A. Zayed, A. Zingone, K. Labadie, J. Ferland, C. Marec, S. Kandels, M. Picheral, C. Dimier, J. Poulain, S. Pisarev, M. Carmichael, S. Pesant, Tara Oceans Coordinators, M. Babin, E. Boss, D. Iudicone, O. Jaillon, S. G. Acinas, H. Ogata, E. Pelletier, L. Stemmann, M. B. Sullivan, S. Sunagawa, L. Bopp, C. de Vargas, L. Karp-Boss, P. Wincker, F. Lombard, C. Bowler, L. Zinger, Global trends in marine plankton diversity across kingdoms of life. Cell 179, 1084–1097.e21 (2019).

129. F. Mahé, N. Henry, C. Berney, J. Poulain, S. Romac, G. Bourdin, H.-J. Ruscheweyh, G. Salazar, C. Belser, Q. Clayssen, B. C. C. Hume, E. Boissin, P. E. Galand, S. Pesant, F. Lombard, E. Armstrong, N. Lang Yona, G. Klinges, R. McMinds, R. Vega Thurber, C. Moulin, S. Agostini, B. Banaigs, E. Boss, C. Bowler, E. Douville, J. M. Flores, D. Forcioli, P. Furla, E. Gilson, S. Reynaud, M. B. Sullivan, O. Thomas, R. Troublé, D. Zoccola, S. Planes, D. Allemand, S. Sunagawa, C. R. Voolstra, P. Wincker, C. de Vargas, rDNA 18S V9 ASV tables (SWARM) for Tara Pacific Expedition, Zenodo (2024); 10.5281/ZENODO.10822169.

