## Supplemental Material for "Diversity, host-specificity, and environmental drivers of the coral reef eukaryome"

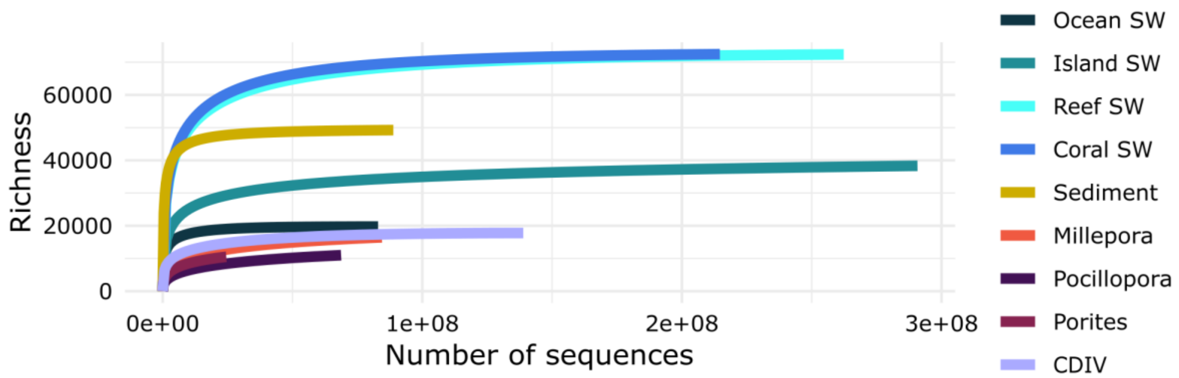


**Supplementary Figure 1.** Rarefaction curves showing the accumulation of amplicon sequence variant (ASV) richness as a function of sequencing depth for each sample type: ocean, island, reef and coral seawater, sediment and the corals *Millepora*, *Pocillopora* and *Porites* from the Tara Pacific expedition.

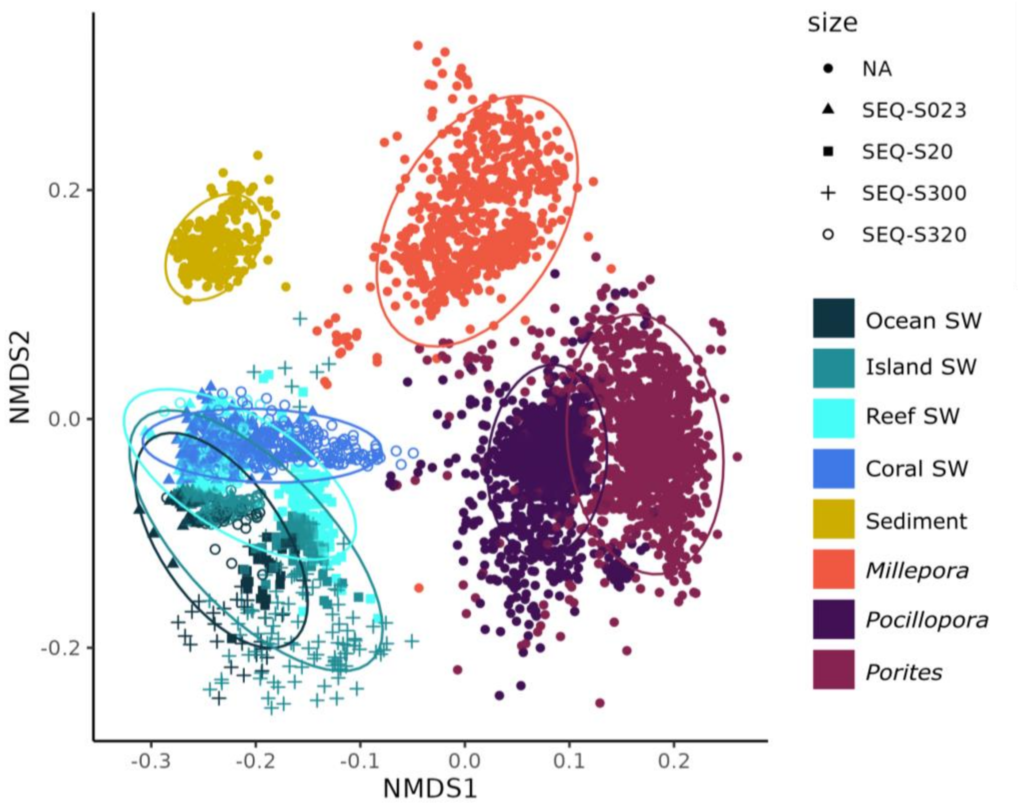


**Supplementary Figure 2.** nMDS of sample types (excluding CDIV) based on Jaccard dissimilarity (stress = 0.14) distances of ASVs, showing the size fractionation. Colours indicate sample types, and shapes represent fraction sizes. Fraction size data were available only for ocean, island and reef seawater samples (see Methods in Lombard et al. 2023).

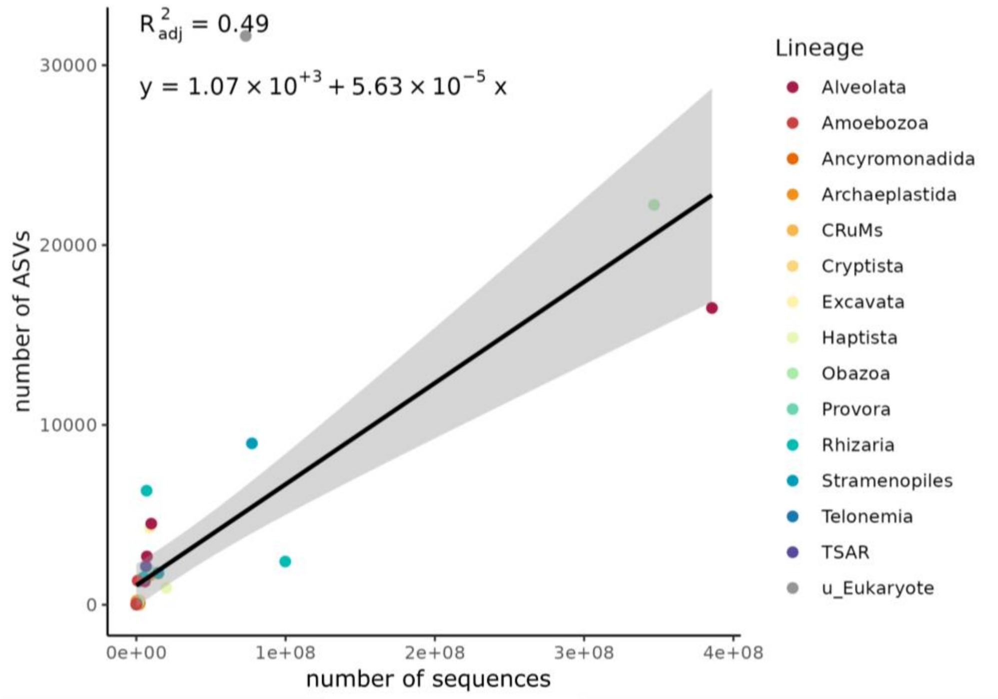


**Supplementary Figure 3.** Relationship between the number of sequences and the number of ASVs for the different major monophyletic groups detected in the *Tara* Pacific Expedition through 18S metabarcoding. The data correspond to Figure 3.

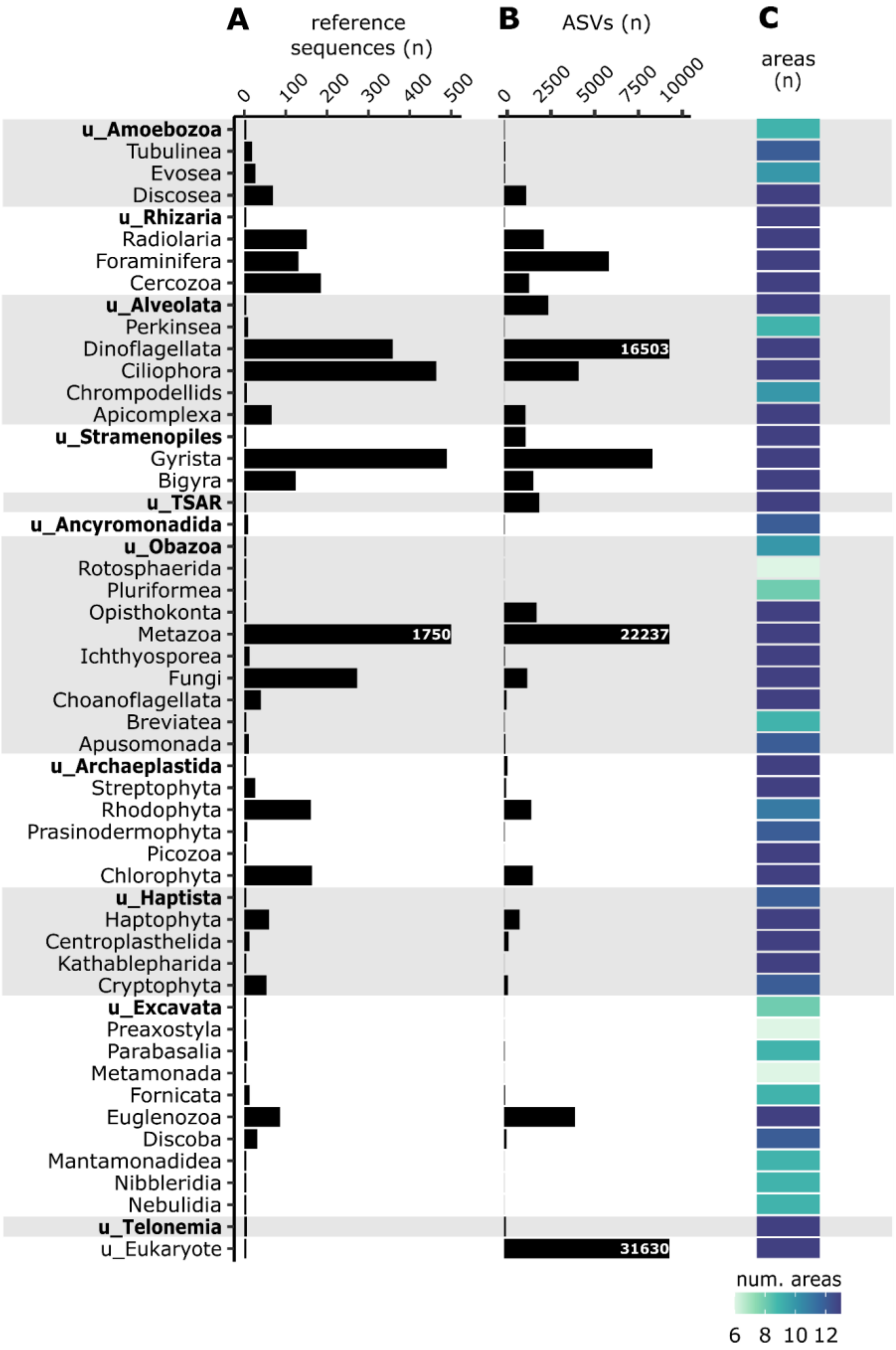


**Supplementary Figure 4. (A)** Number of V9 reference sequences used for taxonomic annotation in each lineage from PR2 v.5.0.0 database (Guillou et al. 2013); bars are labelled when values exceed the axis range. **(B)** Observed richness expressed as the number of ASVs; bars are labelled when values exceed the axis range. **(C)** Number of biogeographic areas where the lineage has been found. Grey shading indicates taxonomic groups belonging to the same major lineage (names in bold). The prefix "u_" indicates unclassified taxa that could only be assigned to higher taxonomic ranks because they lack sufficient sequence similarity to known reference sequences.

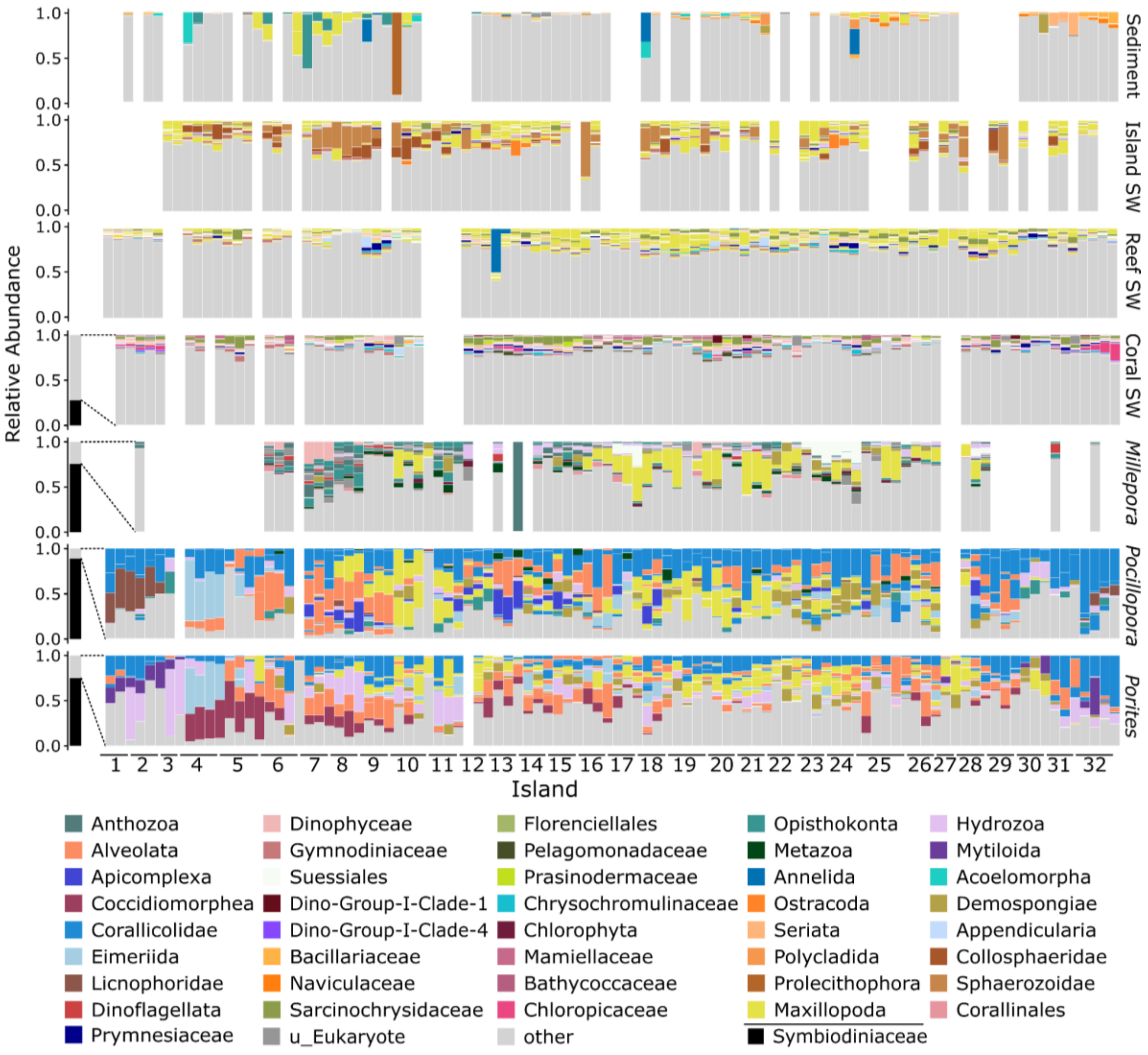


**Supplementary Figure 5.** Relative abundance of microeukaryotic ASVs for each sample type across islands. Each bar represents a site. For clarity, Symbiodiniaceae ASV abundances are shown separately, as they dominate sequence counts in coral samples. Sites where only corals or only the environment were collected have been excluded (102/113 sites kept).

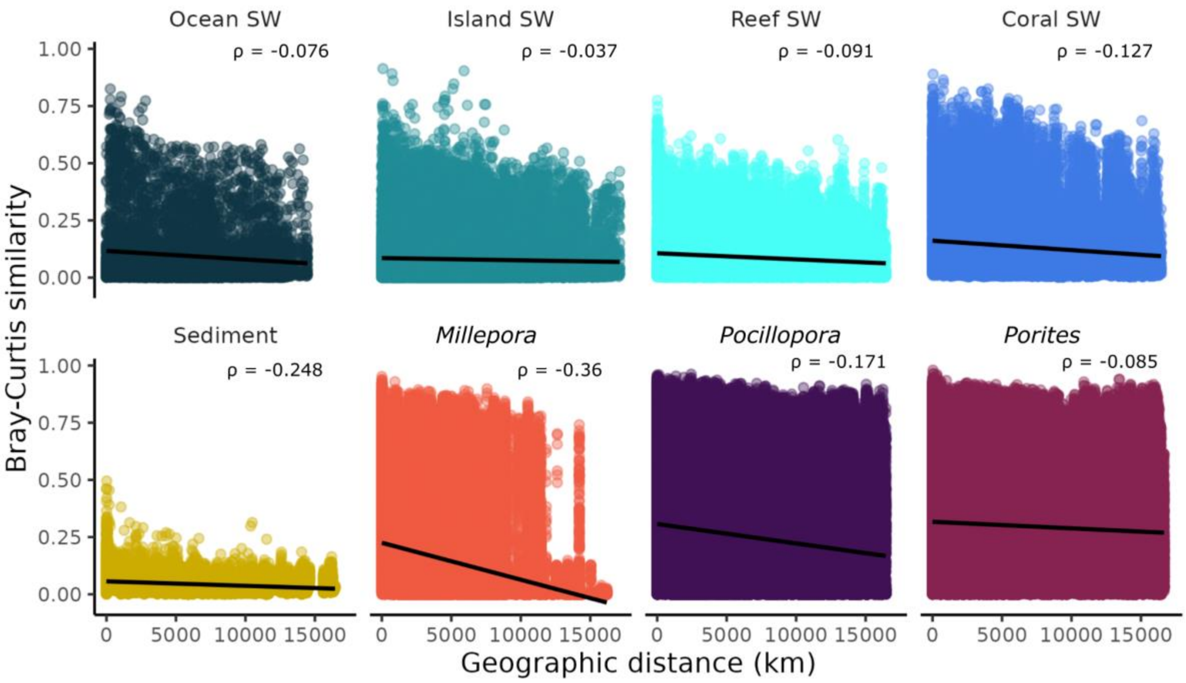


**Supplementary Figure 6.** Decay of community similarity with geographic distance between sites per sample type, based on Bray–Curtis dissimilarity and Haversine pairwise geographic distances.

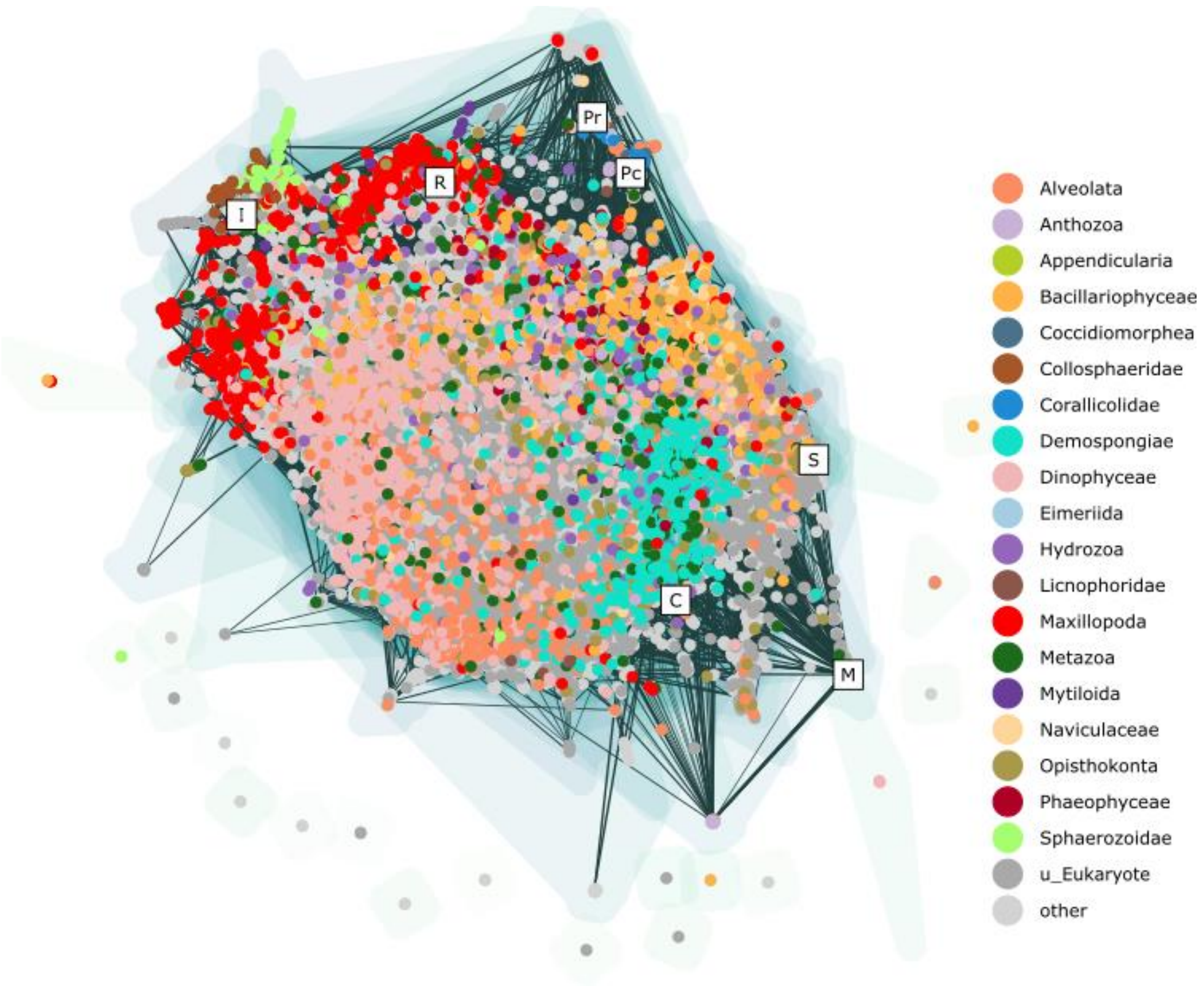


**Supplementary Figure 7.** Alternative visualization of the microeukaryotes network of the Pacific (all) with the centroid of each sample type labelled - *Millepora* (M), *Pocillopora* (Pc) and *Porites* (Pr), Ocean (O), Island (I), Reef (R) and Coral surrounding (C) seawaters, and Sediments (S), and within the corals sampled. The coral communities do not include Symbiodiniaceae. Nodes represent unique ASVs and are colored by the lowest taxonomic rank identification possible. Lines connect nodes with positive associations, and line width is related to the strength of that relationship. Shades represent clusters of ASVs, calculated with Louvain.

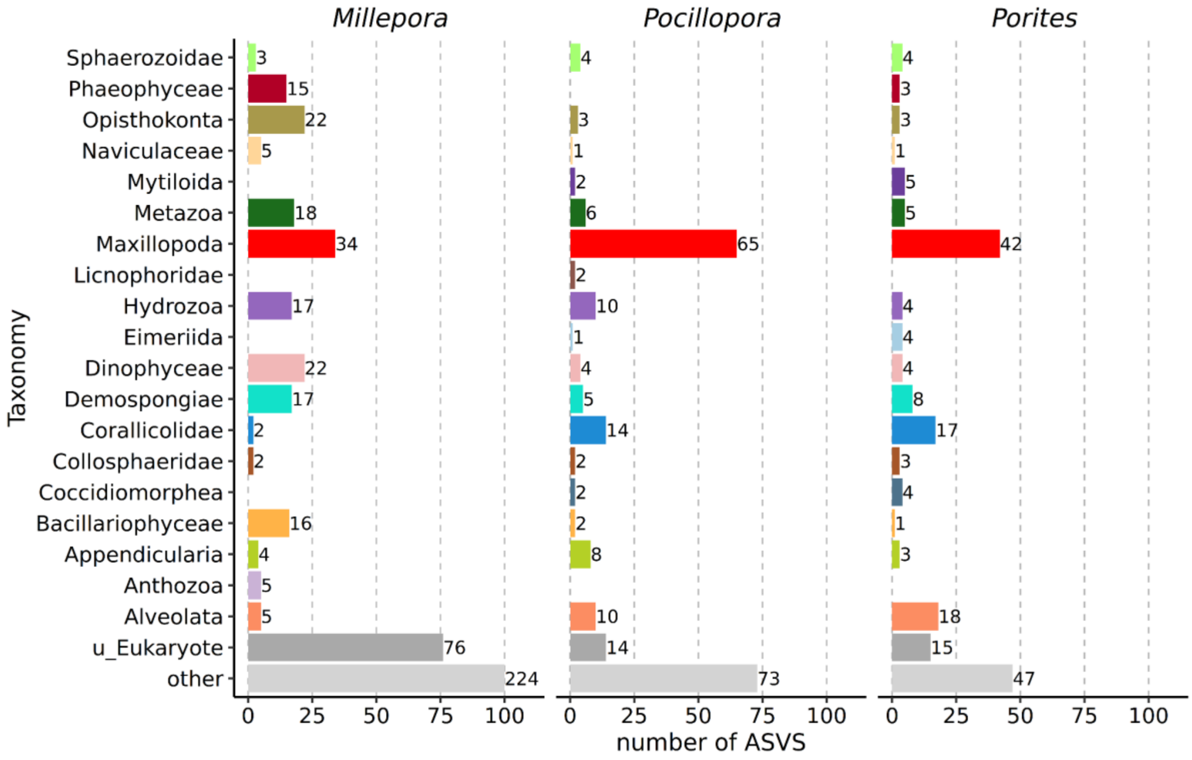


**Supplementary Figure 8.** Number of ASVs per taxonomic category found in the network analysis for *Millepora*, *Porites* and *Pocillopora* morphotypes. Note that in *Millepora*, the category "other" exceeds the x-axis.

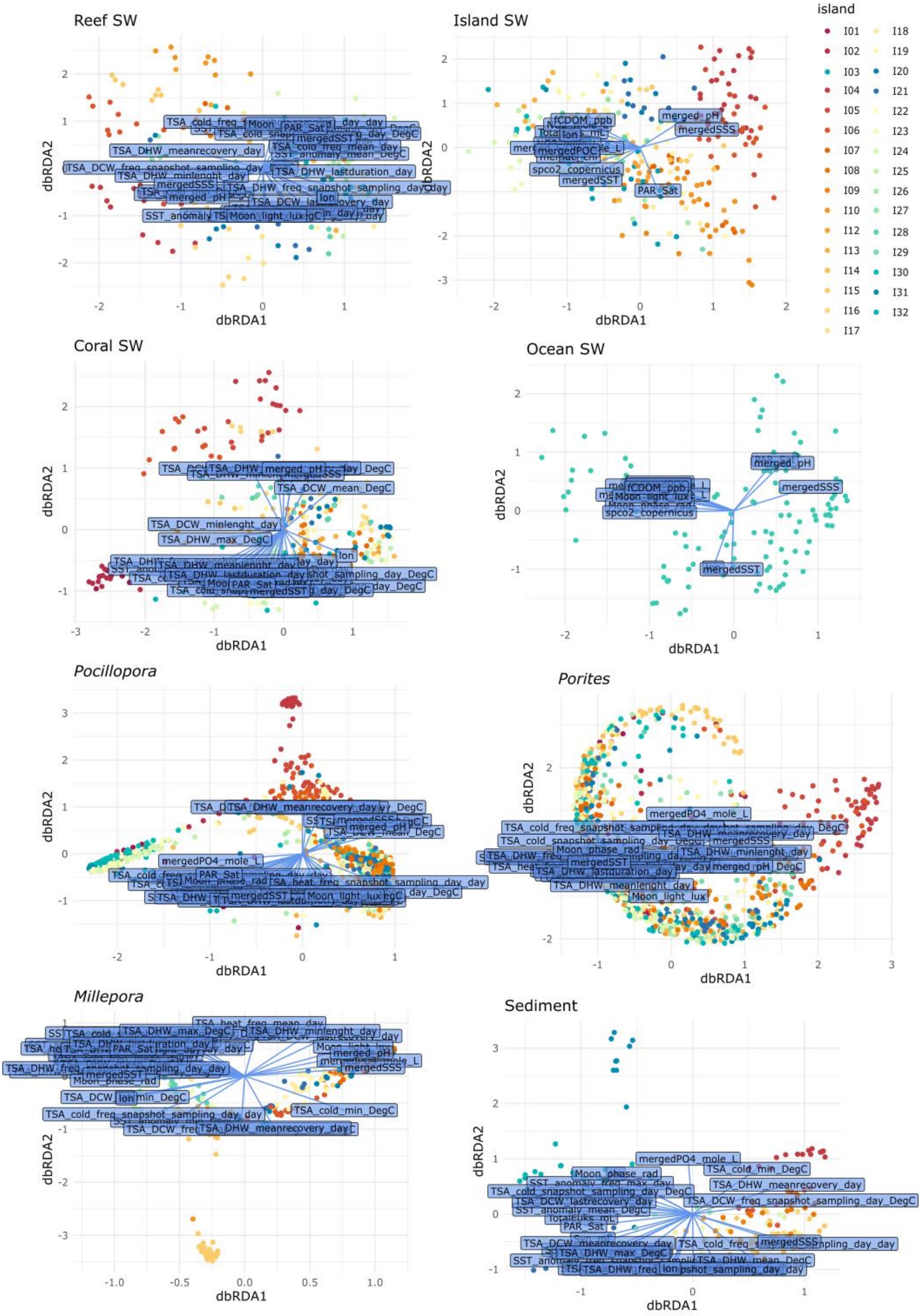


**Supplementary Figure 9:** Redundancy Analysis (RDA) plots for each sample type. Points represent samples and are colored by island.

**Supplementary Table 1.** Filtering steps applied to the raw ASV dataset to retain only target ASVs and samples.

| | **ASV** | | **Samples** | |
|---|---|---|---|---|
| | removed | retained | removed | retained |
| Raw data | - | 262,666 | - | 6,690 |
| Bacteria and Archaea | *-11,646* | *251,020* | *-5* | *6,685* |
| no_hit | *-113,944* | 127,076 | *-2* | 6,683 |
| Skin fungi ASVs | *- 2* | 127,074 | *- 0* | 6,683 |
| Unwanted sample types (fish, aerosol) | *- 191* | 126,883 | *- 34* | 6,449 |
| Atlantic samples | *- 3276* | 123,607 | *- 112* | 6,337 |
| Mammals ASVs | *- 3* | 123,604 | *- 0* | 6,337 |
| Anthozoan ASVs in Scleractinian samples and Coral SW, as well as *Millepora* ASVs in *Millepora* samples | *- 2,372* | 121,232 | *- 0* | 6,337 |

**Supplementary Table 2.** Number of sequences and relative abundances of Anthozoans ASVs in ocean, island and reef seawater (SW) samples.

| **Sample type** | **Number of sequences** | **Relative abundance** |
|---|---:|---:|
| Ocean SW | 10,212 | 0.0001230841 |
| Island SW | 23,8323 | 0.0008196467 |
| Reef SW | 216,1073 | 0.0082376096 |

**Supplementary Table 3.** Number of sequences and relative abundance of sequences that returned no close hit reference sequence (*no_hit*) against the PR2 v5.0.0 dataset (Guillou et al. 2023) across the different sample types included in this study.

| **Sample type** | **Total number of sequences** | ***no_hit* sequences** | ***no_hit* sequences (%)** |
|---|---|---|---|
| Ocean SW | 482.1 $\times 10^6$ | 40.2 $\times 10^6$ | 8.34 |
| Island SW | 347.3 $\times 10^6$ | 32.7 $\times 10^6$ | 9.42 |
| Reef SW | 324.7 $\times 10^6$ | 32.2 $\times 10^6$ | 9.92 |
| Coral surrounding SW | 474.4 $\times 10^6$ | 41.2 $\times 10^6$ | 8.70 |
| Sediment | 163.1 $\times 10^6$ | 41.6 $\times 10^6$ | 25.48 |
| Millepora | 273.5 $\times 10^6$ | 3.1 $\times 10^6$ | 1.14 |
| Pocillopora | 324.4 $\times 10^6$ | 0.2 $\times 10^6$ | 0.07 |
| Porites | 306.8 $\times 10^6$ | 0.4 $\times 10^6$ | 0.14 |
| CDIV | 720.6 $\times 10^6$ | 8.3 $\times 10^6$ | 1.16 |

**Supplementary Table 4.** Full names, descriptions, and original data sources of the environmental variables used in the analysis.

| Variable | Description | Source |
|---|---|---|
| lat | mean latitude of the station\|\|Units:decimal degree\|\|Origin:registery | Bourdin et al. (2022a) |
| lon | mean longitude of the station\|\|Units:decimal degree\|\|Origin:registery | Bourdin et al. (2022a) |
| Syn_mL | Synechococcus concentration\|\|Units:nb.ml-1\|\|Origin:FCM | Bourdin et al. (2022a) |
| TotalEuks_mL | Total eukaryotes concentration\|\|Units:nb.ml-1\|\|Origin:FCM | Bourdin et al. (2022a) |
| merged_chl | Chlorophyll a\|\|Units:mg.m^-3\|\|Origin:HPLC samples/ACS/Satellite/Mercator | Bourdin et al. (2022a) |
| NO2_mole_L | NO2 concentration\|\|Units:µmole/L\|\|Origin:NUT | Bourdin et al. (2022a) |
| NO3_mole_L | NO3 concentration\|\|Units:µmole/L\|\|Origin:NUT | Bourdin et al. (2022a) |
| mergedPO4_mole_L | PO4 concentration\|\|Units:µmole/L\|\|Origin:NUT/Mercator | Bourdin et al. (2022a) |
| merged_SiOH_mole_L | Si(OH)4 concentration\|\|Units:µmole/L\|\|Origin:NUT/Mercator | Bourdin et al. (2022a) |
| Moon_phase_rad | moon phase\|\|Units:radian\|\|Origin:from almanachs using lat/lon+time | Bourdin et al. (2022a) |
| Moon_light_lux | light sent from moon (do not includes position)\|\|Units:lux\|\|Origin:from almanachs using lat/lon+time | Bourdin et al. (2022a) |
| mergedSST | Sea surface temperature\|\|Units:°C\|\|Origin:TSG/Satellite/Mercator | Bourdin et al. (2022a) |
| mergedSSS | Sea surface salinity\|\|Units:psu\|\|Origin:Samples/TSG/Satellite/Mercator | Bourdin et al. (2022a) |
| PAR_Sat | Photosynthtically available radiations\|\|Units:µmol quanta m-2 s-1\|\|Origin:MODIS AQUA sattelite L3 mapped product | Bourdin et al. (2022a) |
| merged_pH | pH \|\|Units:-\|\|Origin:CARB/pH/Mercator | Bourdin et al. (2022a) |
| SST_anomaly_snapshot_sampling_day_DegC | SST anomaly (SST seasonal average) at sampling date (Degree C) | Bourdin et al. (2022b) |
| SST_anomaly_min_DegC | minimum SST anomaly (SST seasonal average) between 2002 and sampling date (Degree C) | Bourdin et al. (2022b) |
| SST_anomaly_max_DegC | maximum SST anomaly (SST seasonal average) between 2002 and sampling date (Degree C) | Bourdin et al. (2022b) |
| SST_anomaly_mean_DegC | average SST anomaly (SST seasonal average) between 2002 and sampling date (Degree C) | Bourdin et al. (2022b) |
| SST_anomaly_freq_snapshot_sampling_day_day | SST anomaly frequency (nb days SST_anomaly >= 1 for the past 52 weeks) at sampling date (days) | Bourdin et al. (2022b) |
| SST_anomaly_freq_max_day | maximum SST anomaly frequency (nb days SST_anomaly >= 1 for the past 52 weeks) between 2002 and sampling date (days) | Bourdin et al. (2022b) |
| TSA_DCW_snapshot_sampling_day_DegC | TSA degree cooling week (sum of previous 12 weeks when TSA_cold <= 1) at sampling date (Degree C) | Bourdin et al. (2022b) |
| TSA_DCW_mean_DegC | average TSA degree cooling week (sum of previous 12 weeks when TSA_cold <= 1) between 2002 and sampling date (Degree C) | Bourdin et al. (2022b) |
| TSA_DCW_freq_snapshot_sampling_day_DegC | TSA degree cooling week frequency (nb days TSA_DCW <= 1 for the past 52 weeks) at sampling date (days) | Bourdin et al. (2022b) |
| TSA_DCW_minlenght_day | minimum TSA degree cooling week event length (nb of consecutive days of TSA_DCW <= 1) between 2002 and sampling date (days) | Bourdin et al. (2022b) |
| TSA_DCW_lastmin_DegC | minimum last event of TSA degree cooling week (Degree C) | Bourdin et al. (2022b) |
| TSA_DCW_meanrecovery_day | average length of recovery period between events of TSA degree cooling week between 2002 and sampling date (days) | Bourdin et al. (2022b) |

| | | |
|---|---|---|
| TSA_DCW_lastrecovery_day | length of last recovery period since the last event of TSA degree cooling week (days) | Bourdin et al. (2022b) |
| TSA_cold_snapshot_sampling_day_DegC | cold Thermal Stress Anomaly at sampling date (Degree C) | Bourdin et al. (2022b) |
| TSA_cold_min_DegC | minimum cold Thermal Stress Anomaly between 2002 and sampling date (Degree C) | Bourdin et al. (2022b) |
| TSA_cold_freq_snapshot_sampling_day_day | cold Thermal Stress Anomaly frequency (nb days TSA_cold <= 1 for the past 52 weeks) at sampling date (days) | Bourdin et al. (2022b) |
| TSA_cold_freq_mean_day | average cold Thermal Stress Anomaly frequency (nb days TSA_cold <= 1 for the past 52 weeks) between 2002 and sampling date (days) | Bourdin et al. (2022b) |
| TSA_heat_freq_snapshot_sampling_day_day | heat Thermal Stress Anomaly frequency (nb days TSA_heat >= 1 for the past 52 weeks) at sampling date (days) | Bourdin et al. (2022b) |
| TSA_heat_freq_mean_day | average heat Thermal Stress Anomaly frequency (nb days TSA_heat >= 1 for the past 52 weeks) between 2002 and sampling date (days) | Bourdin et al. (2022b) |
| TSA_DHW_snapshot_sampling_day_DegC | TSA degree heating week (sum of previous 12 weeks when TSA_heat >= 1) at sampling date (Degree C) | Bourdin et al. (2022b) |
| TSA_DHW_max_DegC | maximum TSA degree heating week (sum of previous 12 weeks when TSA_heat >= 1) between 2002 and sampling date (Degree C) | Bourdin et al. (2022b) |
| TSA_DHW_mean_DegC | average TSA degree heating week (sum of previous 12 weeks when TSA_heat >= 1) between 2002 and sampling date (Degree C) | Bourdin et al. (2022b) |
| TSA_DHW_freq_snapshot_sampling_day_day | TSA degree heating week frequency (nb days TSA_DHW >= 1 for the past 52 weeks) at sampling date (days) | Bourdin et al. (2022b) |
| TSA_DHW_minlenght_day | minimum TSA degree heating week event length (nb of consecutive days of TSA_DHW >= 1) between 2002 and sampling date (days) | Bourdin et al. (2022b) |
| TSA_DHW_meanlenght_day | average TSA degree heating week event length (nb of consecutive days of TSA_DHW >= 1) between 2002 and sampling date (days) | Bourdin et al. (2022b) |
| TSA_DHW_lastduration_day | length of the last event of TSA degree heating week (Degree C) | Bourdin et al. (2022b) |
| TSA_DHW_meanrecovery_day | average length of recovery period between events of TSA degree heating week between 2002 and sampling date (days) | Bourdin et al. (2022b) |

**Supplementary Table 5.** Representative environmental variables selected from groups of correlated variables. For each representative variable, the table lists the correlated variables it represents.

| **Representative variable** | **Correlated variables** |
|---|---|
| SST_anomaly_freq_max_day: maximum SST anomaly frequency (nb days SST_anomaly >= 1 for the past 52 weeks) between 2002 and sampling date (days) | |
| | SST_anomaly_freq_std_day: standard deviation SST anomaly frequency (nb days SST_anomaly >= 1 for the past 52 weeks) between 2002 and sampling date (days) |
| SST_anomaly_min_DegC: minimum SST anomaly (SST - seasonal average) between 2002 and sampling date (Degree C) | |
| | SST_anomaly_freq_mean_day: average SST anomaly frequency (nb days SST_anomaly >= 1 for the past 52 weeks) between 2002 and sampling date (days) |
| | SST_anomaly_std_DegC: standard deviation SST anomaly (SST - seasonal average) between 2002 and sampling date (Degree C) |
| SST_anomaly_snapshot_sampling_day_DegC: SST anomaly (SST - seasonal average) at sampling date (Degree C) | |
| | TSA_heat_max_DegC: maximum heat Thermal Stress Anomaly between 2002 and sampling date (Degree C) |
| TSA_cold_freq_mean_day: average cold Thermal Stress Anomaly frequency (nb days TSA_cold <= -1 for the past 52 weeks) between 2002 and sampling date (days) | |
| | TSA_cold_freq_max_day: maximum cold Thermal Stress Anomaly frequency (nb days TSA_cold <= -1 for the past 52 weeks) between 2002 and sampling date (days) |
| | TSA_cold_freq_std_day: standard deviation cold Thermal Stress Anomaly frequency (nb days TSA_cold <= -1 for the past 52 weeks) between 2002 and sampling date (days) |
| TSA_DCW_lastmin_DegC: maximum last event of TSA degree cooling week (Degree C) | |
| | TSA_DCW_lastsum_DegC: sum last event of TSA degree cooling week (Degree C) |
| TSA_DCW_mean_DegC: average TSA degree cooling week (sum of previous 12 weeks when TSA_cold <= -1) between 2002 and sampling date (Degree C) | |
| | TSA_DCW_min_DegC: minimum TSA degree cooling week (sum of previous 12 weeks when TSA_cold <= -1) between 2002 and sampling date (Degree C) |
| | TSA_DCW_sum_DegC: sum TSA degree cooling week (sum of previous 12 weeks when TSA_cold <= -1) between 2002 and sampling date (Degree C) |
| | TSA_cold_mean_DegC: average cold Thermal Stress Anomaly between 2002 and sampling date (Degree C) |
| | TSA_cold_freq_sum_day: sum cold Thermal Stress Anomaly frequency (nb days TSA_cold <= -1 for the past 52 weeks) between 2002 and sampling date (days) |
| | TSA_cold_std_DegC: standard deviation cold Thermal Stress Anomaly between 2002 and sampling date (Degree C) |
| | SST_std_DegC: standard deviation SST time series between 2002 and sampling date (Degree C) |
| | seasonal_std_DegC: standard deviation of seasonal average of SST between 2002 and sampling date (Degree C) |
| TSA_DCW_meanrecovery_day: average length of recovery period between events of TSA degree cooling week between 2002 and sampling date (days) | |

TSA_DCW_minrecovery_day: minimum length of recovery period between events of TSA degree cooling week between 2002 and sampling date (days)

TSA_DCW_maxrecovery_day: maximum length of recovery period between events of TSA degree cooling week between 2002 and sampling date (days)

mergedSST: Sea surface temperature||Units:°C||Origin:TSG/Satellite/Mercator

TSA_DCW_meanlenght_day: average TSA degree cooling week event length (nb of consecutive days of TSA_DCW <= -1) between 2002 and sampling date (days)

TSA_DCW_lastduration_day: length of the last event of TSA degree cooling week (Degree C)

TSA_DCW_stdlenght_day: standard deviation TSA degree cooling week event length (nb of consecutive days of TSA_DCW <= -1) between 2002 and sampling date (days)

TSA_DCW_freq_sum_day: sum TSA degree cooling week frequency (nb days TSA_DCW <= -1 for the past 52 weeks) between 2002 and sampling date (days)

TSA_DCW_freq_mean_day: average TSA degree cooling week frequency (nb days TSA_DCW <= -1 for the past 52 weeks) between 2002 and sampling date (days)

TSA_DCW_freq_std_day: standard deviation TSA degree cooling week frequency (nb days TSA_DCW <= -1 for the past 52 weeks) between 2002 and sampling date (days)

TSA_DCW_freq_max_DegC: maximum TSA degree cooling week frequency (nb days TSA_DCW <= -1 for the past 52 weeks) between 2002 and sampling date (days)

TSA_DCW_maxlenght_day: maximum TSA degree cooling week event length (nb of consecutive days of TSA_DCW <= -1) between 2002 and sampling date (days)

TSA_DCW_std_DegC: standard deviation TSA degree cooling week (sum of previous 12 weeks when TSA_cold <= -1) between 2002 and sampling date (Degree C)

SST_mean_DegC: average SST time series between 2002 and sampling date (Degree C)

SST_max_DegC: maximum SST time series between 2002 and sampling date (Degree C)

SST_snapshot_sampling_day_DegC: SST interpolated at sampling date (Degree C)

SST_min_DegC: minimum SST time series between 2002 and sampling date (Degree C)

seasonal_mean_DegC: average of seasonal average of SST between 2002 and sampling date (Degree C)

seasonal_min_DegC: minimum seasonal average of SST between 2002 and sampling date (Degree C)

seasonal_max_DegC: maximum seasonal average of SST between 2002 and sampling date (Degree C)

seasonal_snapshot_sampling_day_DegC: seasonal average of SST at sampling date (Degree C)

TSA_heat_snapshot_sampling_day_DegC: Thermal Stress Anomaly at sampling date (Degree C)

TSA_heat_mean_DegC: average heat Thermal Stress Anomaly between 2002 and sampling date (Degree C)

TSA_DHW_meanrecovery_day: average length of recovery period between events of TSA degree heating week between 2002 and sampling date (days)

TSA_DHW_maxrecovery_day: maximum length of recovery period between events of TSA degree heating week between 2002 and sampling date (days)

TSA_DHW_lastrecovery_day: length of last recovery period since the last event of TSA degree heating week (days)

TSA_DHW_minrecovery_day: minimum length of recovery period between events of TSA degree heating week between 2002 and sampling date (days)

TSA_DHW_mean_DegC: average TSA degree heating week (sum of previous 12 weeks when TSA_heat >= 1) between 2002 and sampling date (Degree C)

TSA_DHW_sum_DegC: sum TSA degree heating week (sum of previous 12 weeks when TSA_heat >= 1) between 2002 and sampling date (Degree C)

TSA_DHW_freq_sum_day: sum TSA degree heating week frequency (nb days TSA_DHW >= 1 for the past 52 weeks) between 2002 and sampling date (days)

TSA_DHW_freq_mean_day: average TSA degree heating week frequency (nb days TSA_DHW >= 1 for the past 52 weeks) between 2002 and sampling date (days)

TSA_DHW_stdlenght_day: standard deviation TSA degree heating week event length (nb of consecutive days of TSA_DHW >= 1) between 2002 and sampling date (days)

TSA_DHW_std_DegC: standard deviation TSA degree heating week (sum of previous 12 weeks when TSA_heat >= 1) between 2002 and sampling date (Degree C)

TSA_DHW_freq_max_day: maximum TSA degree heating week frequency (nb days TSA_DHW >= 1 for the past 52 weeks) between 2002 and sampling date (days)

TSA_DHW_maxlenght_day: maximum TSA degree heating week event length (nb of consecutive days of TSA_DHW >= 1) between 2002 and sampling date (days)

TSA_DHW_freq_std_day: standard deviation TSA degree heating week frequency (nb days TSA_DHW >= 1 for the past 52 weeks) between 2002 and sampling date (days)

TSA_DHW_lastduration_day: length of the last event of TSA degree heating week (Degree C)

TSA_DHW_lastsum_DegC: sum last event of TSA degree heating week (Degree C)

TSA_DHW_lastmax_DegC: maximum last event of TSA degree heating week (Degree C)

merged_chl: Chlorophyll a||Units:mg.m^-3||Origin:HPLC samples/ACS/Satellite/Mercator

x19_Butanoyloxyfucoxanthin: 19'-Butanoyloxyfucoxanthin||Units:mg.m^-3||Origin:HPLC

Fucoxanthin: Fucoxanthin||Units:mg.m^-3||Origin:HPLC

Chlorophyll_a: Chlorophyll a + allomers + epimers||Units:mg.m^-3||Origin:HPLC

Diadinoxanthin: Diadinoxanthin||Units:mg.m^-3||Origin:HPLC

CorrectedBact_mL: total bacteria concentration||Units:nb.ml-1||Origin:FCM

phyc_copernicus: Phytoplankton Carbon concentration||Units:mmol m-3||Origin:Copernicus marine service model (reanalysis)

Sum_Carotenes: beta carotene + a-carotene||Units:mg.m^-3||Origin:HPLC

Chlorophyll_c1_c2: sum of chlorophyll c1, c2 and MgDVP||Units:mg.m^-3||Origin:HPLC

Chlorophyll_c3: Chlorophyll c3||Units:mg.m^-3||Origin:HPLC

TotalEuks_mL: Total eukaryotes concentration||Units:nb.ml-1||Origin:FCM

Nano_mL: Nano-eukaryotes concentration||Units:nb.ml-1||Origin:FCM

**Supplementary Table 6.** Overview of the total dataset and each sample type, as well as the raw data. For each category, the number of samples, ASVs, and sequences is reported. The most abundant and most prevalent ASV for each group are also indicated.

| | total | Ocean W | Island W | Reef W | Coral SW | Sediment | *Pocillopora* | *Porites* | *Millepora* | CDIV | raw-data |
|---|---|---|---|---|---|---|---|---|---|---|---|
| samples | 6,337 | 127 | 330 | 281 | 345 | 219 | 1,109 | 1,082 | 652 | 2,192 | 6,690 |
| ASVs | 121,232 | 19,754 | 38,327 | 72,385 | 72,465 | 49,246 | 10,988 | 10,498 | 16,451 | 17,774 | 262,666 |
| total sequences | 1,255 $x10^6$ | 83 $x10^6$ | 291 $x10^6$ | 262 $x10^6$ | 215 $x10^6$ | 89 $x10^6$ | 68 $x10^6$ | 24 $x10^6$ | 84 $x10^6$ | 139 $x10^6$ | 3,417 $x10^6$ |
| min sequences in a sample | 154 | 87,872 | 44,627 | 30,303 | 20653 | 31,760 | 849 | 154 | 2,746 | 308 | 1 |
| max sequences in a sample | 11 $x10^6$ | 8 $x10^6$ | 5 $x10^6$ | 11 $x10^6$ | 5 $x10^6$ | 1 $x10^6$ | 1 $x10^6$ | 1 $x10^6$ | 2 $x10^6$ | 1 $x10^6$ | 12 $x10^6$ |
| max sequences of one ASV | 65 $x10^6$ Symbiodin | 6 $x10^6$ Radiolaria | 16 $x10^6$ Radiolaria | 5 $x10^6$ Crustacean | 32 $x10^6$ u_Symbiodin | 1 $x10^6$ Bacilliario | 30 $x10^6$ u_Symbio | 13 $x10^6$ Cladocopium | 27 $x10^6$ u_Sym | 21 $x10^6$ Cladocopium | - |
| most prevalent ASV | 5661 Cladocopium | 127 u_Hydrozoa | 330 u_Hydrozoa | 281 Pocillopora | 345 u_Gymnodin. | 219 u_Bacilliario | 1046 Cladocopium | 1081 Cladocopium | 650 u_Anthozoa | 2174 Cladocopium | - |

**Supplementary Table 7.** Wilcox's pairwise comparison of Richness (A) and Pielou's Evenness Index (B) between different sample types. P-value adjusted with False Discovery Rate (FDR) method; shaded in grey are the non-significant values (p>0.05).

**A.** Richness

| | CDIV | Coral SW | Island SW | *Millepora* | Ocean SW | *Pocillopora* | *Porites* | Reef SW |
|---|---|---|---|---|---|---|---|---|
| Coral SW | 7.85E-195 | . | . | . | . | . | . | . |
| Island SW | 2.50E-186 | 5.01E-22 | . | . | . | . | . | . |
| *Millepora* | 1.08E-26 | 1.68E-147 | 1.20E-134 | . | . | . | . | . |
| Ocean SW | 1.03E-79 | 5.55E-11 | 0.442343 | 5.67E-67 | . | . | . | . |
| *Pocillopora* | 1.37E-31 | 2.28E-172 | 2.23E-165 | 1.28E-59 | 2.24E-75 | . | . | . |
| *Porites* | 1.68E-141 | 1.45E-171 | 6.79E-165 | 3.06E-120 | 1.99E-75 | 2.67E-43 | . | . |
| Reef SW | 1.00E-163 | 0.998227 | 3.31E-20 | 1.89E-128 | 1.73E-11 | 2.00E-147 | 7.67E-147 | . |
| Sediment | 2.03E-131 | 2.55E-29 | 0.320417 | 4.76E-107 | 0.342475 | 1.08E-120 | 2.35E-120 | 3.52E-20 |

**B.** Pielou's Evenness Index

| | CDIV | Coral SW | Island SW | *Millepora* | Ocean SW | *Pocillopora* | *Porites* | Reef SW |
|---|---|---|---|---|---|---|---|---|
| Coral SW | 5.53E-121 | . | . | . | . | . | . | . |
| Island SW | 9.25E-76 | 0.011485 | . | . | . | . | . | . |
| *Millepora* | 0.444953 | 5.15E-96 | 1.32E-60 | . | . | . | . | . |
| Ocean SW | 4.62E-35 | 0.444953 | 0.436814 | 5.66E-32 | . | . | . | . |
| *Pocillopora* | 3.86E-05 | 4.91E-104 | 8.74E-62 | 0.004303 | 2.10E-30 | . | . | . |
| *Porites* | 1.28E-23 | 9.48E-86 | 1.63E-45 | 1.76E-14 | 1.92E-23 | 2.83E-08 | . | . |
| Reef SW | 7.54E-130 | 0.213681 | 3.04E-05 | 4.56E-108 | 0.042989 | 6.45E-119 | 4.28E-101 | . |
| Sediment | 2.22E-86 | 0.968281 | 0.003319 | 1.12E-74 | 0.230582 | 7.88E-79 | 5.82E-65 | 0.292643 |

**Supplementary Table 8.** PERMANOVA (A) and pairwise PERMANOVA (B) results based on the Jaccard distance matrix, and betadispersion tests (B) with pairwise Tukey's HSD comparisons (D), for eukaryotic community composition across sample types.

### A. PERMANOVA

Permutation test for Adonis under reduced model
Number of permutations: 999

| | Df | SumOfSqs | R2 | F | Pr(>F) |
|---|---|---|---|---|---|
| Model | 7 | 297.71 | 0.15502 | 108.42 | 0.001 |
| Residuals | 4137 | 1622.75 | 0.84498 | | |
| Total | 4144 | 1920.46 | 1.00000 | | |

### B. Pairwise PERMANOVA

| pairs | Df | SumsOfSqs | F.Model | R2 | p.value | p.adjusted |
|---|---|---|---|---|---|---|
| Reef SW vs Island SW | 1 | 3.039856 | 6.667514 | 0.01083 | 0.001 | 0.028 |
| Reef SW vs Coral SW | 1 | 6.283304 | 14.29091 | 0.022389 | 0.001 | 0.028 |
| Reef SW vs Ocean SW | 1 | 2.622476 | 5.798697 | 0.014081 | 0.001 | 0.028 |
| Reef SW vs Pocillopora | 1 | 41.08843 | 108.0301 | 0.072211 | 0.001 | 0.028 |
| Reef SW vs Porites | 1 | 46.87572 | 129.9172 | 0.087139 | 0.001 | 0.028 |
| Reef SW vs Millepora | 1 | 25.45055 | 59.36137 | 0.059939 | 0.001 | 0.028 |
| Reef SW vs Sediment | 1 | 8.252782 | 17.79358 | 0.034497 | 0.001 | 0.028 |
| Island SW vs Coral SW | 1 | 10.88799 | 24.58798 | 0.035247 | 0.001 | 0.028 |
| Island SW vs Ocean SW | 1 | 1.299975 | 2.853628 | 0.006233 | 0.001 | 0.028 |
| Island SW vs Pocillopora | 1 | 46.09262 | 120.0833 | 0.077121 | 0.001 | 0.028 |
| Island SW vs Porites | 1 | 52.2568 | 143.148 | 0.092166 | 0.001 | 0.028 |
| Island SW vs Millepora | 1 | 27.54815 | 63.85067 | 0.061168 | 0.001 | 0.028 |
| Island SW vs Sediment | 1 | 8.759398 | 18.81655 | 0.033256 | 0.001 | 0.028 |
| Coral SW vs Ocean SW | 1 | 6.609635 | 15.22993 | 0.031387 | 0.001 | 0.028 |
| Coral SW vs Pocillopora | 1 | 43.59248 | 115.446 | 0.073652 | 0.001 | 0.028 |
| Coral SW vs Porites | 1 | 57.79826 | 161.0449 | 0.101539 | 0.001 | 0.028 |
| Coral SW vs Millepora | 1 | 33.94635 | 80.51311 | 0.07486 | 0.001 | 0.028 |
| Coral SW vs Sediment | 1 | 12.09219 | 27.03871 | 0.045903 | 0.001 | 0.028 |
| Ocean SW vs Pocillopora | 1 | 21.9133 | 59.10146 | 0.045705 | 0.001 | 0.028 |
| Ocean SW vs Porites | 1 | 24.72643 | 70.94323 | 0.055514 | 0.001 | 0.028 |
| Ocean SW vs Millepora | 1 | 14.5143 | 34.30173 | 0.04228 | 0.001 | 0.028 |
| Ocean SW vs Sediment | 1 | 6.255412 | 13.39921 | 0.037491 | 0.001 | 0.028 |
| Pocillopora vs Porites | 1 | 99.26494 | 284.0456 | 0.114857 | 0.001 | 0.028 |
| Pocillopora vs Millepora | 1 | 86.5393 | 226.1592 | 0.113925 | 0.001 | 0.028 |
| Pocillopora vs Sediment | 1 | 29.82701 | 78.33407 | 0.05578 | 0.001 | 0.028 |
| Porites vs Millepora | 1 | 97.43635 | 265.2507 | 0.132808 | 0.001 | 0.028 |
| Porites vs Sediment | 1 | 34.27113 | 95.11484 | 0.068226 | 0.001 | 0.028 |
| Millepora vs Sediment | 1 | 17.46554 | 40.35104 | 0.044373 | 0.001 | 0.028 |

### C. Betadispersion tests

PERMDISP2 model:

Homogeneity of multivariate dispersions

No. of Positive Eigenvalues: 2999
No. of Negative Eigenvalues: 1145

Average distance to median:

| Coral SW | Island SW | Millepora | Ocean SW | Pocillopora | Porites | Reef SW | Sediment |
|---|---|---|---|---|---|---|---|
| 0.6526 | 0.6753 | 0.6437 | 0.6666 | 0.5896 | 0.5706 | 0.6719 | 0.689 |

Eigenvalues for PCoA axes:
(Showing 8 of 4144 eigenvalues)

| PCoA1 | PCoA2 | PCoA3 | PCoA4 | PCoA5 | PCoA6 | PCoA7 | PCoA8 |
|---|---|---|---|---|---|---|---|
| 173.11 | 137.89 | 79.26 | 65.55 | 57.94 | 55.92 | 47.09 | 31.95 |

Permutation test for homogeneity of multivariate dispersions:

Number of permutations: 999
Response: Distances

| | Df | SumOfSqs | Mean Sq | F | N. Perm | Pr(>F) |
|---|---|---|---|---|---|---|
| Groups | 7 | 7.478 | 1.06828 | 127.6 | 999 | 0.001 |
| Residuals | 4137 | 34.635 | 0.00837 | | | |

**D.** Pairwise Tukey's HSD test on betadispersion. Significant p-values indicated in bold.

| **pair** | **null.value** | **estimate** | **conf.low** | **conf.high** | **adj.p.value** |
|---|---|---|---|---|---|
| ISRFa-Coral SW | 0 | 0.022658 | 0.001294 | 0.044023 | **0.028585** |
| MilleporaCoral SW | 0 | -0.00892 | -0.02739 | 0.009549 | 0.826212 |
| MilleporaIsland SW | 0 | -0.03158 | -0.05033 | -0.01284 | **9.38E-06** |
| OSRF-Coral SW | 0 | 0.014002 | -0.0148 | 0.0428 | 0.821261 |
| OSRF-Island SW | 0 | -0.00866 | -0.03763 | 0.020317 | 0.985579 |
| OSRF-Millepora | 0 | 0.022925 | -0.00399 | 0.049837 | 0.162252 |
| PocilloporaCoral SW | 0 | -0.06304 | -0.08015 | -0.04594 | **3.28E-08** |
| PocilloporaIsland SW | 0 | -0.0857 | -0.1031 | -0.0683 | **3.28E-08** |
| PocilloporaMillepora | 0 | -0.05412 | -0.06781 | -0.04043 | **3.28E-08** |
| PocilloporaOcean SW | 0 | -0.07705 | -0.10304 | -0.05105 | **3.28E-08** |
| PoritesCoral SW | 0 | -0.08208 | -0.09923 | -0.06492 | **3.28E-08** |
| PoritesIsland SW | 0 | -0.10473 | -0.12218 | -0.08729 | **3.28E-08** |
| PoritesMillepora | 0 | -0.07315 | -0.08691 | -0.0594 | **3.28E-08** |
| PoritesOcean SW | 0 | -0.09608 | -0.1221 | -0.07005 | **3.28E-08** |
| PoritesPocillopora | 0 | -0.01903 | -0.03089 | -0.00718 | **3.21E-05** |
| SedimentCoral SW | 0 | 0.036349 | 0.012377 | 0.060322 | **1.19E-04** |
| SedimentIsland SW | 0 | 0.013691 | -0.01049 | 0.037874 | 0.676307 |
| SedimentMillepora | 0 | 0.045272 | 0.023602 | 0.066942 | **4.01E-08** |
| SedimentOcean SW | 0 | 0.022347 | -0.0086 | 0.053294 | 0.358046 |

| | | | | | |
|---|---|---|---|---|---|
| SedimentPocillopora | 0 | 0.099392 | 0.078875 | 0.119909 | **3.28E-08** |
| SedimentPorites | 0 | 0.118425 | 0.097865 | 0.138984 | **3.28E-08** |
| SedimentReef SW | 0 | 0.017081 | -0.00793 | 0.042091 | 0.434048 |
| SSRF-Coral SW | 0 | 0.019269 | -0.00303 | 0.041565 | 0.148655 |
| SSRF-Island SW | 0 | -0.00339 | -0.02591 | 0.019133 | 0.999817 |
| SSRF-Millepora | 0 | 0.028191 | 0.008391 | 0.047991 | **4.29E-04** |
| SSRF-Ocean SW | 0 | 0.005267 | -0.0244 | 0.034934 | 0.999452 |
| SSRF-Pocillopora | 0 | 0.082312 | 0.063781 | 0.100842 | **3.28E-08** |
| SSRF-Porites | 0 | 0.101344 | 0.082766 | 0.119921 | **3.28E-08** |
